# A Spiking Neural Network Model for Category Learning

**DOI:** 10.1101/2020.01.23.916593

**Authors:** Laxmi R. Iyer, Arindam Basu

## Abstract

The creation of useful categories from data is an important cognitive ability, and from the extensive research on categorization, it is now known that the brain has distinct systems for category learning. In this paper, we present the first spiking neural network (SNN) model of human category learning. Here categories are combinations of features - such categories are observed in the prefrontal cortex (PFC). The system follows an architecture commonly used to model the cortex - features are arranged in a topological 2D grid with short range excitatory connectivity and long range inhibitory connectivity - however, here, this architecture is used differently from other models to model higher level cognition. Earlier we presented an artificial neural network (ANN) model of category learning; however, here, a simpler model was adequate, as desired functionality emerges from the SNN dynamics. We identified the objectives that had to be fulfilled for the model to achieve the desired functionality, and performed a design space exploration (DSE) to identify the parameter range in which each of the objectives was fulfilled, and the parameter range for which the system exhibits good performance. Finally, we compared triphasic STDP (a variant of spike time dependant plasticity (STDP)) with standard STDP and observed that triphasic STDP exhibited quicker convergence.

## 1. Introduction

Creating patterns and structure out of the large amount of sensory data the brain receives – thus learning categories which can be used to make further inferences is a major aspect of cognition. The mechanisms underlying this ability are still not fully understood, and it is being extensively studied by psychologists and neuroscientists.

It is now known that category learning involves several distinct systems in the brain [1]. A description of these is given in our earlier paper [2], however, we repeat them here for completeness. Early theories described rule based categorization where categories are combinations of simple features and can be explicitly stated in terms of a rule. Rule based categorization has been associated with the prefrontal cortex (PFC) and anterior cingulate (ACC). Models of rule based categorization include COVIS [3] and RULEX [4]. Prototype category learning defines categories in terms of a prototype around which instances of the category are distributed. This form of categorization is associated with the occipital lobe [5]. Most models of category learning, including popular models such as ALCOVE [6] and SUSTAIN [7] model prototype category learning. Finally, exemplar based category learning models how humans are able to handle exceptions and model category members explicitly. This form of category learning is associated with the medial temporal lobe (MTL), and there is evidence that sometimes the PFC and the MTL compete during categorization tasks. The model described in our paper falls under rule-based category learning.

We model categories as combinations of several features. There is ample evidence for complex visual categories which are a combination of simple features. Tsunoda et. al. [8, 9] applied imaging techniques to study spatial patterns of activation in the inferotemporal cortex (ITC) by various complex objects, and simplified forms of them. Objects are represented by activations of combination of columns each responding to a specific visual feature of the object. This result has been replicated by many others ([10, 11, 12, 13, 14], etc). However, human imaging studies [5] have found that although ITC is sensitive to perceptual features of stimuli, only the prefrontal cortex (PFC) represents the boundary between actual categories or crucial conjunctions between features. It was seen that PFC neurons conveyed which rule was in effect independent of the cue that signaled the rule and the behavioral response. It is known that the PFC receives extensive sensory information - visual, somatosensory, and auditory information from the occipital, temporal, and parietal cortices, and limbic areas. It has feedback connections to other brain areas, and motor outputs [15]. There is also extensive interconnectivity within different PFC areas so information from different modalities can be assimilated [15]. Hence, the PFC is well situated to encode rule based categories which are a conjunction of various features from different modalities.

We implement the model as a two dimensional map consisting of neurons representing features. Neurons within a small radius are excitatorily connected to one another, and outside that radius, neurons have inhibitory connections. A one dimensional attractor with this property is known as line attractor, and with periodic boundary conditions, this structure is known as a ring attractor. Extending this attractor to two dimensions, it is known as a torus attractor [16]. Mikkulainen [17] has given biological evidence for this connectivity in the cortex - the cortex’ excitatory connections are patchy, and although many long range synapses are excitatory, long range connections are overall inhibitory in nature. It is known that this construct is used to model head direction (HD) in the hippocampus - the excitatory connections activate nearby cells, while the inhibitory connections force cells further away to be silent, resulting in a ‘bump’ of excitatory activity on the torus [16]. This construct has been extensively used by Mikkulainen et. al. [18] to model the visual cortex. They have modelled many functions in the visual cortex including the formation of orientation columns, formation of ocular dominance, [18] object segmentation and binding [19]. This model has also been used by others to model the cortex including [20].

However, these are models of the lower sensory modalities and do not model higher level and more abstract category formation. Ritter [21] argues that the same model constructs for lower sensory modalities (i.e. laterally connected two dimensional spatial maps) can also explain the formation of more abstract maps that are used for motor control and semantic meanings of words. Many types of feature selective cells in different cortical and non-cortical regions were discovered, indicating that complex stimuli are represented by ‘special feature combinations’ and this may be a may be a major encoding strategy in the brain [21].

In many of these models, neurons which are in close proximity self-organize to represent similar features. However, in our model, the features that a neuron represent remains fixed as this is necessary for the creation of categories with very disparate features, and to allow the same feature to be a part of very different categories.

Mikkulainen [21] gives an extensive analysis of the benefits of lateral connections - a few that are applicable to our system are given as follows:

- Lateral connections establish local cooperation and global competition among neurons across large areas, thus making self organization possible.
- They store information for feature binding and grouping
- They form a substrate for encoding memories as attractors in the cortical network.
- Lateral excitatory and inhibitory connections serve different roles in the cortex. As a result, there are two competing mechanisms that prevent one attractor from being strong, and utilize more completely, different mechanisms present in the system (This would be further discussed in the final section of this paper).

Many of these phenomena will be observed later on in this paper.

It is generally known that the brain uses similar operating principles for different phenomena, and Ritter [21] provides evidence that spatial maps are actually used for modeling semantic and motor information. Our model is an implementation of Ritter’s theory - a well-known model that is established biologically, and used to model lower sensory processes will be used, in this paper, to model a higher order phenomenon - the learning of complex categories based on combinations of simple features.

The motivation for the system described in this paper is as follows:

- Although there are many models of category learning in the brain, very few are implemented using neural networks.
- Although there are spiking neural network (SNN) models that perform pattern classification, the model described in this paper is the first SNN model that implements category learning in the brain.
- Although there are several SNN attractor models that utilize 2D attractor networks (especially in vision) none of them are used to model a higher-order cognitive task such as category learning.

Earlier, we had created a model of category learning using artificial neural networks (ANNs). In this paper we show an implementation of the model using spiking neural networks (SNNs). Spiking neural networks (SNNs) are the third generation neural networks and more biologically realistic than their predecessors (i.e. ANNs). Unlike ANN units whose outputs are typically between 0 and 1, spiking neurons incorporate individual spikes, thereby using spatiotemporal patterns to transmit information the way real neurons do. Neuron membrane voltages and synaptic current pulses are modelled explicitly in SNNs. We noted that when modelling the system with SNNs, a simpler model can be used to achieve the desired functionality, as the function arises from the natural dynamics of the system. This is an important step in banishing the homunculus. The emphasis of this paper and its difference from the previous paper on our ANN system is summarized as follows.

- The current system is implemented using SNNs while the previous system was implemented using ANNs.
- The ANN model required certain features to be added just so the functionality would be achieved; however, in the SNN model, the desired functionality emerged from the dynamics of the system. Hence a simpler model was adequate.
  – During the presentation of each concept, the ANN model required recurrent inhibition to start low and increase slowly to prevent all units from immediately becoming inactive. Also the input activation had to be high in the beginning, and was slowly decreased. However, in the SNN model, decaying input EPSC currents achieved both - the influence of input currents decayed over time, and this naturally led to greater influence of recurrent inhibitory currents.
  – In the ANN system, threshold adaptation of neurons was required to prevent a previously well-learnt attractor from being activated for partial stimuli, and therefore creating a new attractor that overlapped with the previous one. In the SNN system, as only excitatory weights are learnt, excitatory recurrent current depends on weights, while inhibitory recurrent current does not. Hence, increasing the decay time constant of inhibitory current increases inhibitory currents in the system, and biases competition towards regions that are not necessarily learned. This prevents the same attractor from being active for irrelevant or partial stimuli.
- The SNN system is analyzed via design space explorations, and the parameter range in which certain objectives that are necessary for proper functionality are met, is identified and the reasons for the same examined. The parameter range in which the system performs well is also identified, and reasons for the same examined.
- The SNN system employs triphasic STDP (T-STDP) [22], a variant of the typically used biphasic STDP [23, 24, 25, 26, 27]. A comparison of results between T-STDP and B-STDP (i.e. standard biphasic STDP) is performed.

The paper is organized as follows: the following section is the Background. Following this is a description of the problem and the features of the ANN category learning model. The category learning framework and problem we are trying to solve is identical to that of the previous ANN system. Therefore, everything described in this section applies to both the SNN and ANN models and has been described earlier, however, it is described here for completeness and in some cases added clarity compared to the previous paper. For the rest of the paper, the words ‘the system’ refers to the SNN system. The next section has a description of the SNN implementation of the system. Following this, we delineate the objectives that the SNN system has to fulfill in order to achieve the desired functionality. Finally we have results which are divided into two subsections, firstly, the design space exploration that explores the parameter range for which each of the above objectives are fulfilled, and for which the system yields good results. Secondly, a comparison of biphasic (standard) STDP and triphasic STDP (the STDP variant used in the previous section) ensues.

## 2. ANN Category Learning Model

The description of this section is based on our previous ANN model [2], and is presented here for completeness. Since the SNN system is based on the same framework, the description applies to the SNN model as well.

### 2.1. Features of the Model

Categories are useful when they do not merely capture the statistical regularities in the data, but are based on task requirements [7]. Studies show that people form categories which are useful for particular tasks, and given extensive exposure to a new task, they will form categories for it. However, the categories thus formed are used generally across different tasks. Therefore, our category learning model postulates that the learning of categories is task-specific and its usage is task-independent. Therefore the features of the system are as follows:

1. Extraction of relevant feature spaces for categorization: After training on a number of concepts, the system should be able to recognize a subset of frequently co-occuring features.
2. Context dependent learning of categories: The system picks out only feature combinations that occur frequently within a context and ignores other combinations and regularities that occur across contexts.

### 2.2. Problem Formulation

In literature, the words ‘category’ and ‘concept’ have been used interchangeably, the word ‘context’ is used to describe a task situation, and the word ‘feature’ usually refers to the simple attributes of objects. To avoid ambiguity, in this paper we use the terms ‘category’, ‘concept’, ‘feature’ and ‘context’ as follows (also illustrated in Table 1).

**Table 1:**
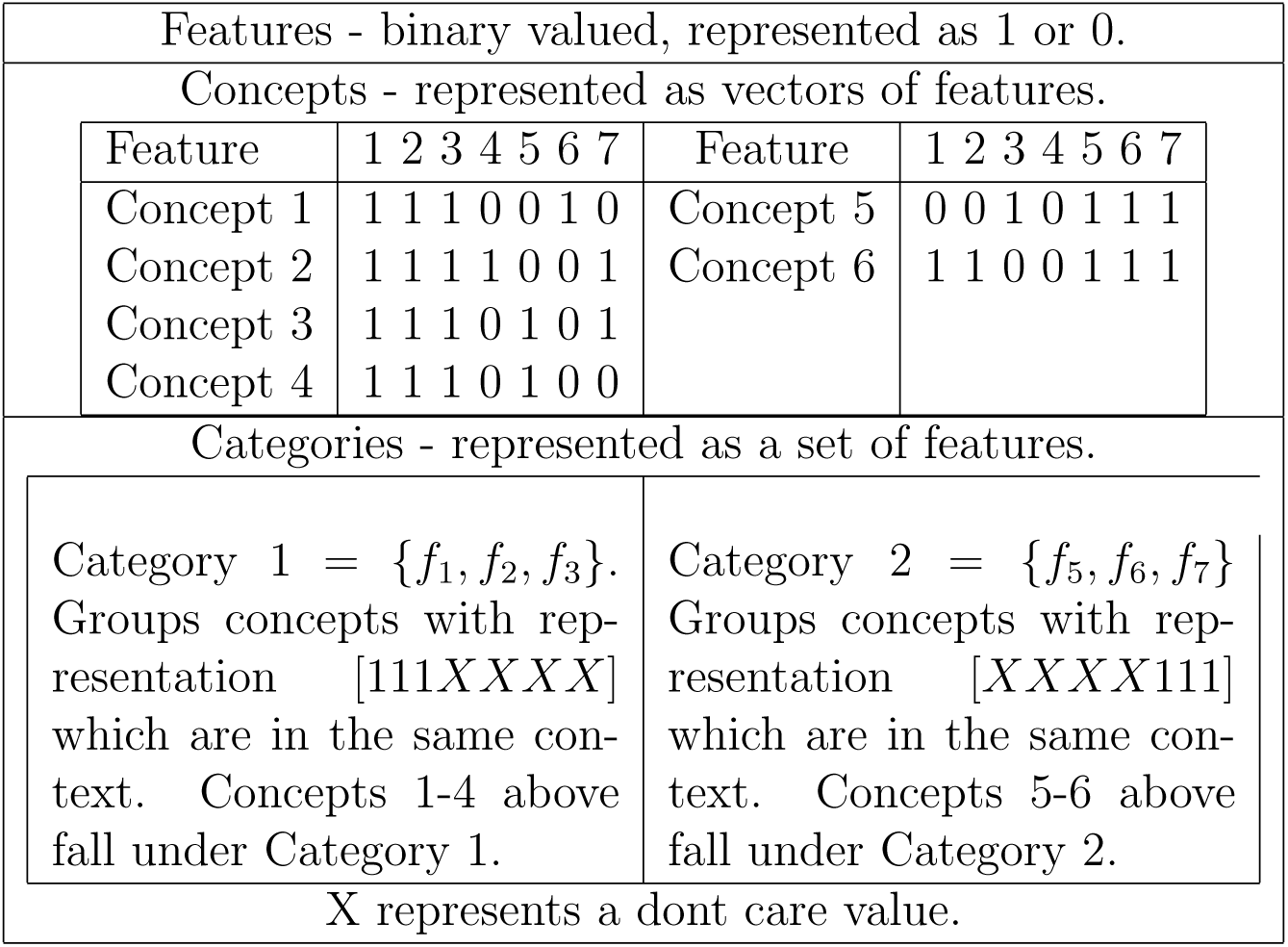
System definitions.

| Features - binary valued, represented as 1 or 0. |  |  |  |  |  |  |  |  |  |  |  |  |  |  |  |
| --- | --- | --- | --- | --- | --- | --- | --- | --- | --- | --- | --- | --- | --- | --- | --- |
| Concepts - represented as vectors of features. |  |  |  |  |  |  |  |  |  |  |  |  |  |  |  |
| Feature | 1 | 2 | 3 | 4 | 5 | 6 | 7 | Feature | 1 | 2 | 3 | 4 | 5 | 6 | 7 |
| Concept 1 | 1 | 1 | 1 | 0 | 0 | 1 | 0 | Concept 5 | 0 | 0 | 1 | 0 | 1 | 1 | 1 |
| Concept 2 | 1 | 1 | 1 | 1 | 0 | 0 | 1 | Concept 6 | 1 | 1 | 0 | 0 | 1 | 1 | 1 |
| Concept 3 | 1 | 1 | 1 | 0 | 1 | 0 | 1 |  |  |  |  |  |  |  |  |
| Concept 4 | 1 | 1 | 1 | 0 | 1 | 0 | 0 |  |  |  |  |  |  |  |  |
| Categories - represented as a set of features. |  |  |  |  |  |  |  |  |  |  |  |  |  |  |  |
| Category 1 = {f<sub>1</sub>, f<sub>2</sub>, f<sub>3</sub>}.<br>Groups concepts with representation [111XXXX] which are in the same context. Concepts 1-4 above fall under Category 1. |  |  |  |  |  |  |  | Category 2 = {f<sub>5</sub>, f<sub>6</sub>, f<sub>7</sub>}.<br>Groups concepts with representation [XXXX111] which are in the same context. Concepts 5-6 above fall under Category 2. |  |  |  |  |  |  |  |
| X represents a dont care value. |  |  |  |  |  |  |  |  |  |  |  |  |  |  |  |

**Table 2:** Simulation Parameters.

|  |  |
| --- | --- |
| Leaky IF Parameters<br>$V_{leak} = -7$ mV<br>$R = 10^5$ Ohms<br>$\tau_M = 30$ ms<br>$V_{th} = -6$ mV<br>$V_{reset} = -8$ mV<br>$v_{ref} = 2$ ms<br>Simulation Parameters<br>$dt = 1$ ms<br>Concept cycle length = 0.5 s<br>$t_1 = 60$ ms<br>$t_2 = 220$ ms<br>$t_3 = 380$ ms<br>Feature Grid Parameters<br>$r = 6$<br>No. of features = 25<br>No. of neurons representing a<br>feature on the grid = 100<br>Size of network:<br>No. of inputs = 25<br>No. of neurons on the grid = 2500 | Synapse Parameters<br>$\tau_{IX} = 60$ ms<br>$\tau_{IN} = 300$ ms<br>$\tau_{RX} = 5$ ms<br>$\tau_{RN} = DSE \text{ Parameter}$<br>$A^{IX} = DSE \text{ Parameter}$<br>$A^{IN} = 1500$ nA<br>$A^{RX} = 15$ nA<br>$A^{RN} = 0.5$ nA<br>STDP Parameters<br>Triphasic STDP<br>$\eta = \frac{\tau_{wd}}{dt} 0.001$<br>$\alpha = 30$ ms<br>Biphasic STDP<br>$A_+ = 3.9$<br>$A_- = 1.5$<br>$\tau_+ = 7$ ms<br>$\tau_- = 262$ ms<br>Weight Decay<br>$W_{st} = 0.7$<br>$\tau_{wd} = 7$ s<br>Frequency Detection<br>$\tau_\xi = 20$ ms<br>$\hat{\xi} = 1.5$ |

1. A *feature* is a simple attribute of an object. Features may indicate the type of the concept, for e.g. ‘food’, ‘tool’, ‘animal’, or a description of the concept, for e.g. ‘red’, ‘green’, ‘long’, ‘square’, etc. Distinct features represent mutually exclusive attributes [28]. Features have a binary value, 1 indicates the presence of a feature and 0 indicates the absence of a feature. Features are the fundamental units in this system. We denote the number of features in our system by *N*_*f*_.
2. A *concept* is a complex object that consists of a specific set of features and not others. Each concept is represented as a binary vector of features [*f*_1_, *f*_2_, *f*_3_, …*f*_*Nf*_] where *f*_*m*_ = 1 if feature *m* is present and *f*_*n*_ = 0 if feature *n* is absent. Concepts are exemplars that are presented to the system during training. The system is trained on *N*_*c*_ concepts.
3. A *context* represents a task situation, and it is expected that within a context concepts will share the same salient features. So in this paper, we define context as a time period where the system experiences concepts with the same sets of co-occurring features repeatedly. A change in the context is denoted by a sudden change in all the salient features (for e.g. if we suddenly go from home to the university we expect to experience different features altogether.)
4. If *m* concepts within a context contain the same *k* features, then the *k* features are said to be frequently co-occurring within the context. These *k* features forms a *category*. So the category can be represented as a set of features {*f*_1_, *f*_2_, …*f*_*k*_}, and groups together concepts of the same context wherein all the features in the set are present, regardless of whether other features are present or absent.

These terms are clearly illustrated in Table 1.

The system experiences several task contexts during each of which a sequence of concepts are presented to the system. Each concept is presented to the system as a vector of features. The goal of the system is to *group concepts within the same context that share a co-occurring set of features (known as salient features)* as a category.

## 3. System Description

The system has one layer, the feature layer, which is a distributed representation of the features. The system functionality is shown in figure 1.

**Figure 1:**
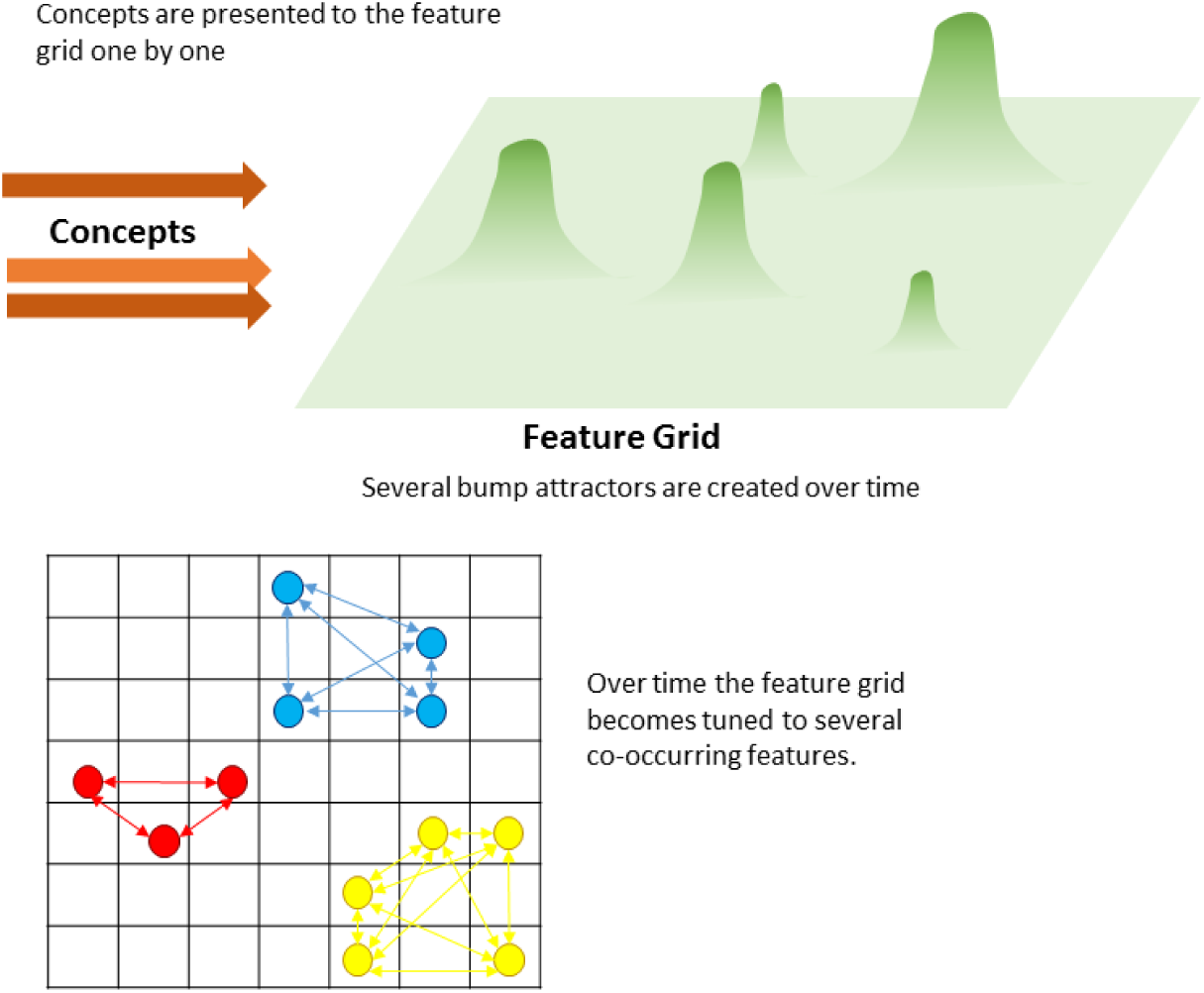
System Functionality - as the system is trained on concepts, the feature grid slowly develops localized attractors each of which consists of a strongly connected set of features.

Each concept is presented to the feature layer as input currents. The feature layer also has lateral excitatory and inhibitory connections. The interplay between input and recurrent currents leads to the formation of strong localized attractors that represent feature correlations.

The dynamics of the layer is described in this section.

### 3.1. Initialization of Feature Layer

The model has a two dimensional layer of neurons each representing a feature. Each feature is represented by several neurons in the feature layer: the function *µ*(*i*) maps neuron *i* to the feature *f*_*k*_ that it represents, and *µ* is a many to one function - one feature represented by many neurons.

Each neuron *i* in the feature layer has two dimensional coordinates *i* = (*x*_*i*_, *y*_*i*_). The feature layer is assumed to have periodic boundary conditions to avoid edge effects. Any neuron *j* is connected to *i* with excitatory connections if the Euclidean distance between them (*d*_*ij*_) is below a value *r* which is a system parameter. The neurons are distributed randomly in the feature grid, in such a way that the distance between each feature is maximized, so many features will fall into a local area in the grid. This is done as follows:

1. Calculate distances *d*_*ij*_ for every pair of neurons in the feature grid.
2. Initialize two variables: first, *available neurons* list is initialized to the neuron IDs of all the neurons in the feature grid. Second, *feature assignment map* is an array of length equal to the number of neurons on the feature grid with all values initialized to 0.
3. For each feature *F*:
  a. Choose a neuron (say *A*) randomly from the *available neurons* list, and assign it to to the current feature *F* - i.e. element *A* in the *feature assignment map* is assigned value *F*.
  b. Remove those neurons within less than distance *r*_1_ from the assigned neuron, from the *available neurons* list. Here we set *r*_1_ to *r*.
  c. Continue till all the instances of a feature have been assigned, or the *available neuron* list is empty.
  d. If there are more instances of a feature to be assigned, and the *available neurons* list is empty, assign the rest of the neurons randomly to neurons that have not been assigned yet.
  e. Reinitialize the *available neurons* list to be equal to list of neurons not currently assigned to any feature. Repeat from step 3.

We use the quantity 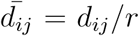, a scaled distance between two neurons to calculate initial weights. Initial excitatory weights between neurons are set according to the distance between them as follows.

The excitatory weight from unit *j* to unit *i* in the feature layer is calculated as:

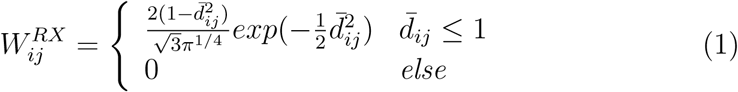

Inhibitory weights in the feature layer 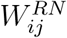 are 0 if 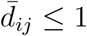 else, they are −1. Therefore *W* ^*RX*^ ∈ [0, 1], *W* ^*RN*^ ∈ {0, −1}.

Excitatory weights are modified during learning while inhibitory weights are not.

### 3.2. Feature Layer Dynamics

The input and recurrent connections are shown in figure 2.

**Figure 2:**
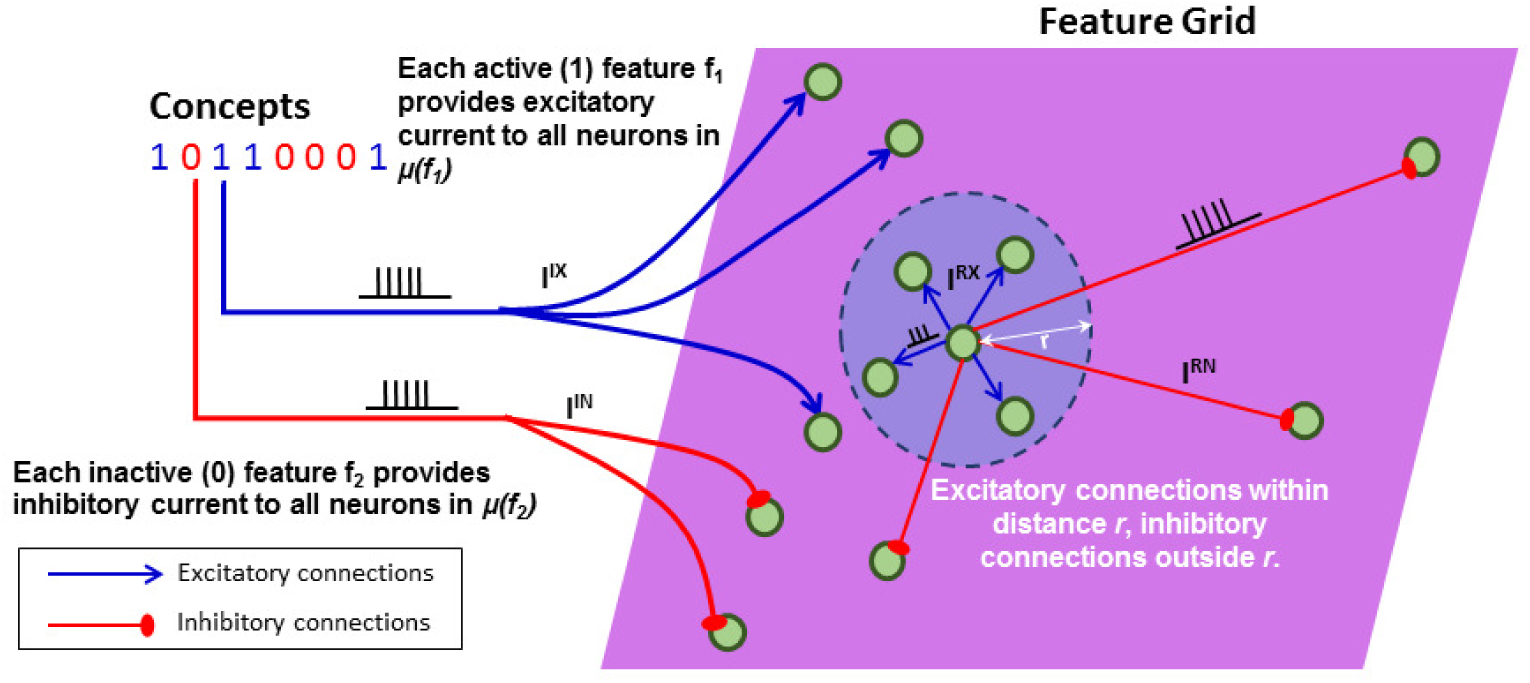
System input and recurrent connections.

**Figure 3:**
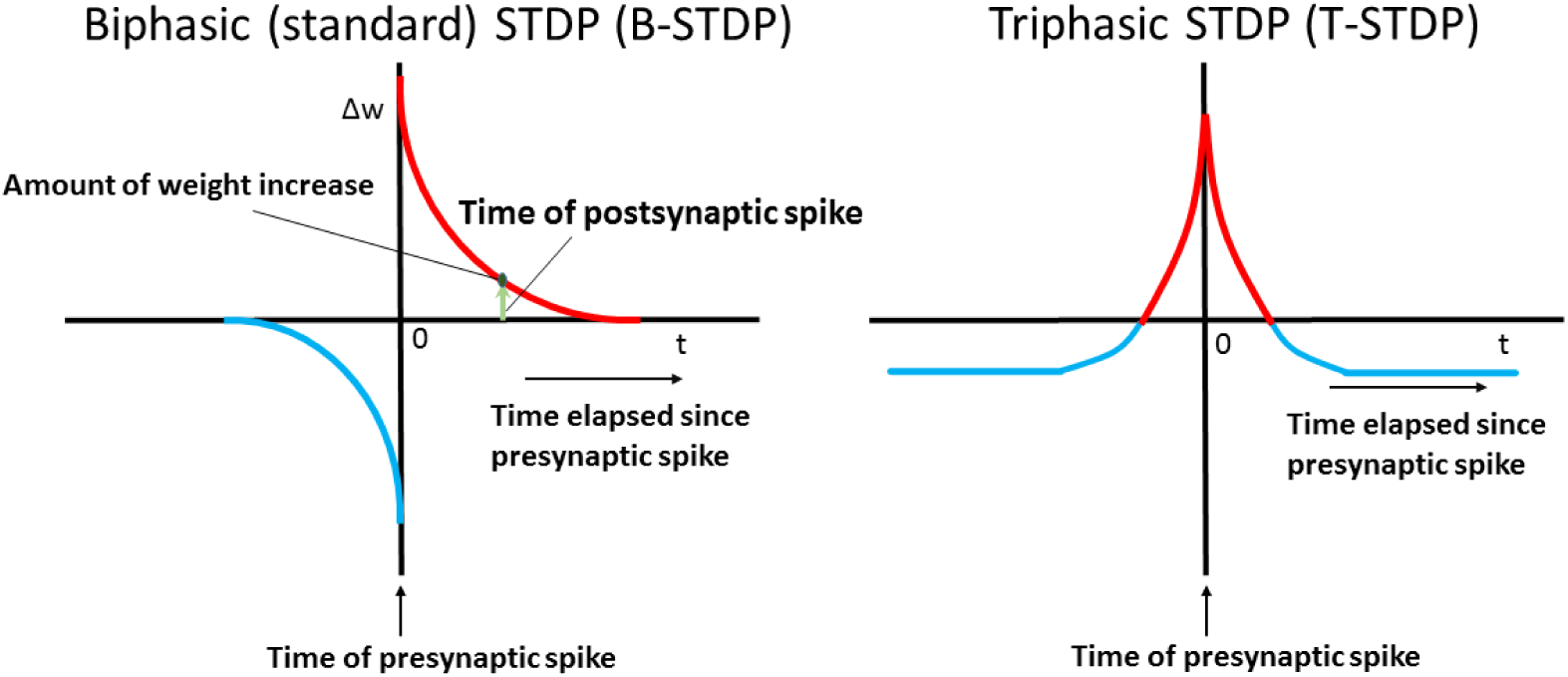
Triphasic (T-STDP) and biphasic (B-STDP) learning rules. While T-STDP is symmetric and does not depend on the order of firing, but only the time difference between the pre- and post-synaptic spikes, B-STDP features potentiation when post- is followed by pre-synaptic spike, and depression when pre- is followed by post-synaptic spike.

**Figure 4:**
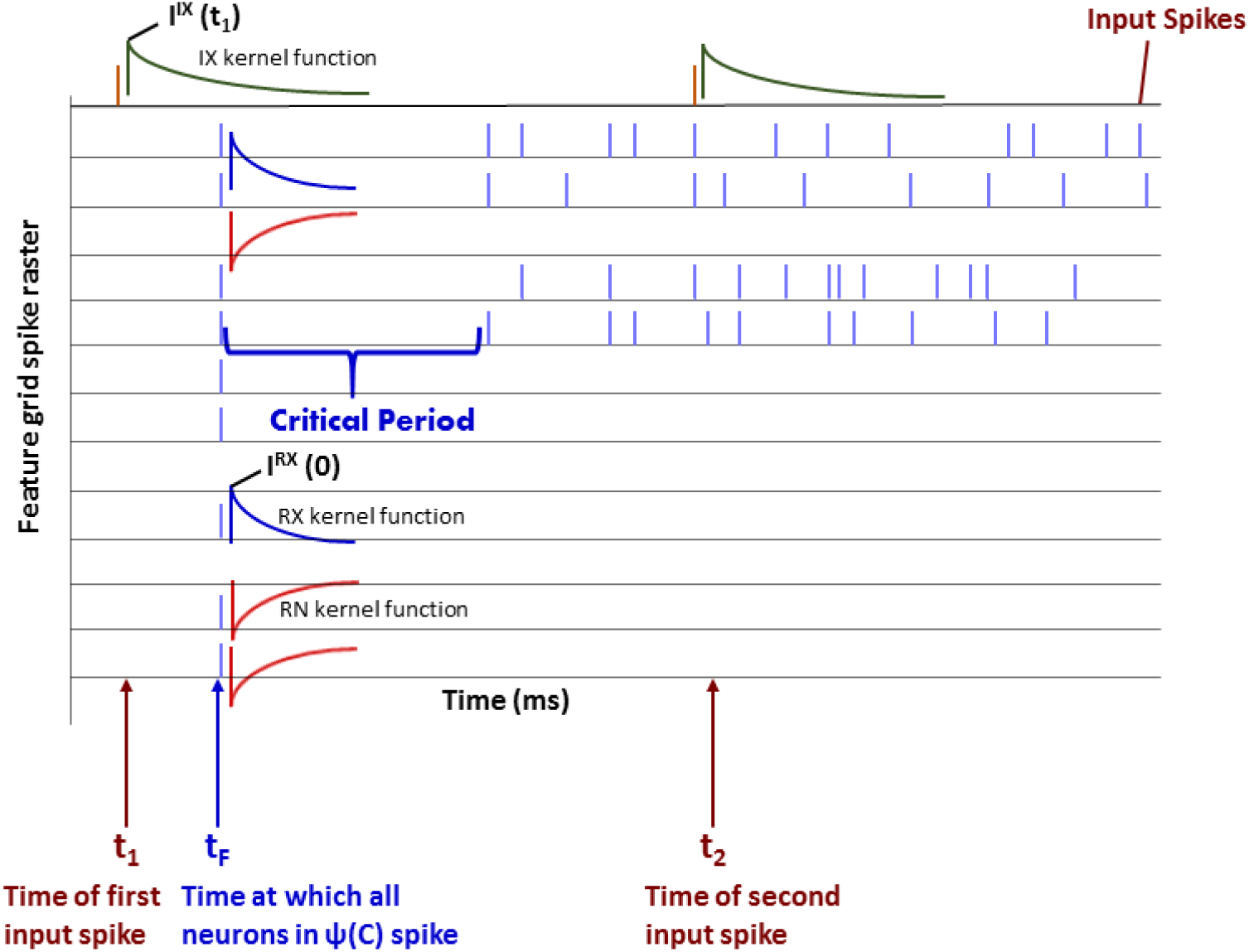
Basis of the simplified model.

When a concept is presented to the feature layer, the corresponding spike trains are 500 ms long - this is the duration of concept presentation which we term a ‘concept cycle’. Towards the end of the concept cycle, the system settles to an attractor. As mentioned earlier, each concept is represented as a binary *N*_*f*_ -dimensional vector of features. The excitatory weights 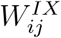 between input *I* and the feature grid *F* denote the connections between each input feature *f*_*i*_ and each neuron *j* on the feature layer that represents it. So for neuron 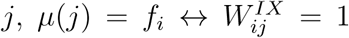 are the inhibitory binary weights and are the negation of 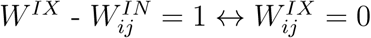 and vice versa. Therefore *W* ^*IX*^, *W* ^*IN*^ ∈ {0, 1} (see figure 2).

During the presentation of a concept *C, C* (which is a binary vector of features) is converted to two *N*_*f*_-dimensional vectors, *C*_*E*_ which is equal to *C* and *C*_*I*_, which is the one’s complement of *C*. Spikes are generated according to the following template:

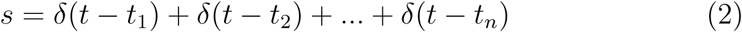

where 0 *< t*_1_ *< t*_2_ *<* … *< t*_*n*_ *<* 500. In these simulations, *n* = 3.

For every element 1 in *C*_*E*_, a spike train is generated according to the template above. Similarly for every element 0 in *C*_*I*_ a spike train is generated according to the template above.

Therefore, there will be two sets of *N*_*f*_ spike trains that will be input into the feature layer, one providing excitatory currents and one providing inhibitory currents into the feature units. If a feature *f*_1_ is 1 in concept *C*, then all neurons {*i*|*µ*(*i*) = *f*_1_} will receive excitatory current with amplitude *A*^*IX*^. If another feature *f*_2_ is 0 in concept *C*, then all neurons {*i*|*µ*(*i*) = *f*_2_} will get inhibitory current with amplitude |*A*^*IN*^|.

|*A*^*IN*^| is far greater than |*A*^*IX*^| and the amplitudes of recurrent currents (which are described below). As a result, neurons that represent features that are not present in the current concept receive a strong inhibitory current, and other currents are not able to overcome them. As a result, the only neurons in the feature layer that participate in the feature layer dynamics during the current concept cycle are those in the set Ψ(*C*) which is defined as follows:

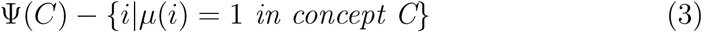

The feature layer is modeled by integrate and fire (IF) neurons [29, 30, 31, 32, 33, 34, 35], etc. and the membrane voltage *V* of neuron *i* at time *t* is given as follows:

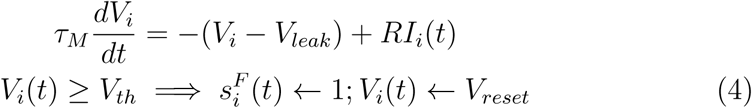

where *τ*_*M*_ is the membrane time constant, *V*_*th*_ is the threshold voltage and *V*_*leak*_ is the leak voltage. *I*_*i*_(*t*) is the total current that is input into neuron *i*. 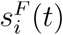 is the spike train of neuron *i* and is 1 during the time neuron *i* fires a spike and 0 otherwise. After a spike is fired, the membrane voltage of the neuron is set to *V*_*reset*_ and the neuron is refractory for the absolute refractory period *v*_*ref*_, before it can fire again.

Each neuron *i* in the feature layer receives four currents (see 2):

1. Input excitatory currents - these are the excitatory currents from the input feature. All variables pertaining to input excitatory currents will hence be denoted with the superscript *IX*.
2. Input inhibitory currents - these are the inhibitory currents from the input features. Relevant variables will be denoted with the superscript *IN*.
3. Recurrent excitatory currents - these are recurrent excitatory currents from the neurons that have excitatory connections with the neuron. Relevant variables will be denoted with the superscript *RX*.
4. Recurrent inhibitory currents - these are recurrent inhibitory connections from the neurons that have inhibitory currents with the neuron. Relevant variables will be denoted with the superscript *RN*.

The total current into each neuron *i* is the summation of the input and recurrent currents, as follows:

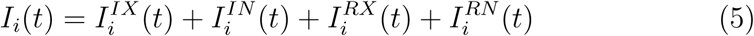

where 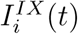 is the input excitatory current, 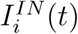 is the input inhibitory current, 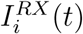 is the recurrent excitatory current, and 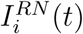 is the recurrent inhibitory current. Each of these currents are calculated as follows:

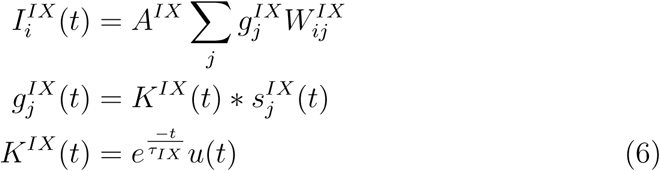

where *u* is the Heaviside function. Similarly, *I*^*IN*^, *I*^*RX*^ and *I*^*RN*^ are calculated using the weights 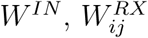 and 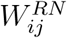 respectively. 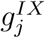 is the *IX* input kernel function. 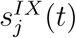 is the *IX* spike train corresponding to feature *j*. Similar equations follow for 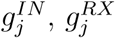 and 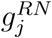, using time constants *τ*_*IN*_, *τ*_*RX*_ and *τ*_*RN*_, for *IN, RX* and *RN* current respectively, using spike trains 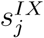 and 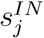 for *IX* and *IN* currents respectively, and 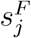 for recurrent (*RX* and *RN*) currents. 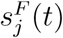 is the spike train of neuron *j* in the feature layer.

The *I*^*IX*^ current provides initial activity to the feature layer, and serves as a ‘booster’ current to ensure that activity in the feature layer remains - else inhibitory currents might deactivate all neurons in the feature layer. Each unit receives *I*^*RX*^ from neurons in its local neighborhood, and *I*^*RN*^ currents from all other neurons. This induces a competition between different local neighborhoods (or ‘regions’) of activity till finally activity persists in one region alone. Neurons within a region spike strongly, and these represent features of the presented concept.

### 3.3. Learning

Excitatory recurrent weights 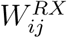 are modified according to the equation below. There is no modification to any other weights.

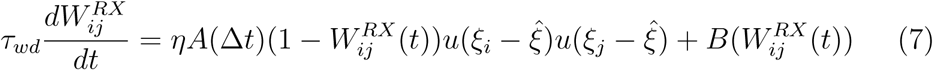

where each of the components are described below. The (1 − *W*) term in the above equation induces a soft bound on the weights, causing them to be in between 0 and 1. Any neuron *i* can have its synaptic weights modified only if its firing rate crosses a threshold - this is to avoid weight increases in neurons that seldom fire. This is done with the internal variable *ξ*_*i*_, and *u* is the Heaviside step function. We tested the system on T-STDP as well as B-STDP - *A*(Δ*t*) has different equations for both the STDP variants. Finally the weights are subject to decay and this is captured by the function 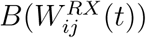.

The equation for *ξ*_*i*_ as follows:

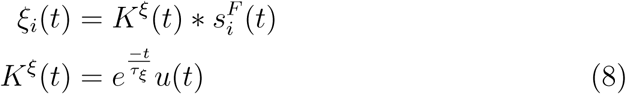

where *τ*_*ξ*_ is the time constant of the variable *ξ*_*i*_ and *u* is the Heaviside function. If 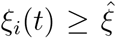 then neuron *i* is eligible to have its weights modified from time *t* onwards.

#### 3.3.1. Triphasic STDP

The neurons learn with the triphasic STDP (T-STDP) learning rule [22], which is a variant of spike time dependant plasticity (STDP). This learning rule was chosen due to the symmetric nature of the learning between neurons - the increase in weight between neuron *i* and *j* is the same as the increase in weights between neuron *j* and *i*. This symmetricity is useful because both neuron *i* and *j* should reinforce each other in a reciprocal manner to sustain attractors that contain frequently co-occuring features. In later sections, we shall see that compared to standard (biphasic) STDP, triphasic STDP has a shorter training time. The equations for learning the weights are as follows:

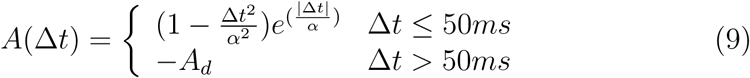

where −*A*_*d*_ is the evaluation of the first part of equation 9 at *t* = 50 ms, and is a small negative value.

#### 3.3.2. Biphasic STDP

We also trained the system with the standard biphasic STDP training rule (B-STDP) [23, 24, 25, 26, 27] and compared the performance with T-STDP training rule. The equations used are:

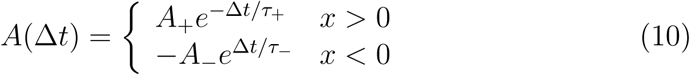

To ensure that both training rules are comparable, the parameters *A*_+_, *A*_−_, *τ*_+_ and *τ*_−_ are set such that integrating the function above over all *t* (area under the curve of the function) yields the same result as the area under the curve of function in equation 9. Also, note that learning rate *η* is the same in both the cases.

#### 3.3.3. Weight Decay

Finally weight decay is described below:

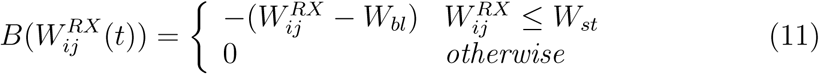

where *τ*_*wd*_ is the weight decay time constant, *W*_*bl*_ are the baseline weights, or the initial weights in the system, and *W*_*st*_ is the stabilization threshold for weights [36]. So all weights that are below the stabilization threshold decay towards their baseline weights.

Neurons get activated throughout the concept cycle, and if they fire with a high rate, they get their weights modified. But at the end of the concept cycle, activity is confined to only one region, and with units firing very rapidly within this region. They fire rapidly because of three reasons:

1. They receive no (or very little) inhibitory current, as only neighboring units are active, and very few neurons outside its excitatory receptive field are active.
2. They receive strong excitatory current because its neighbors are also firing rapidly.
3. As they are firing, their weights are getting strengthened, causing them to receive more excitatory current, and to increase their firing rates even further.

As weights are strengthened, similar input tends to activate the same region, due to strong recurrent currents (due to increased recurrent weights in this region). If certain features tend to co-occur, they repeatedly get activated, and weights between them become stronger, forming a strong localized attractor over time. If other features occur only sporadically, their weight increase during one concept cycle is followed by weight decay during the subsequent concept cycles. So the attractor contains only those features that tend to frequently co-occur.

As defined earlier, a context is a period of time where concepts are expected to share some similar features. So if concepts that have frequently co-occurring features within a context, they are expected to occur closely in time. The similar features, as shown earlier, will form an attractor. On the other hand, if concepts that share several features are not in the same context (and therefore, they will not occur one after the other, but will be temporally separated) then weight increases for those feature sets are followed by weight decay, never allowing them to form an attractor.

### 3.4. Simplified Model

In order to quickly verify if the model meets certain criteria for desirable functionality (described in a later section) and predict the outcome of the design space explorations (explained below) as well as explain the phenomena observed, we devised a simplified model, which is a set of equations that can explain the outcome of the model quickly (a comparison of the speed of the simplified model and the simulations is given at a later section) in different parameter settings.

During the presentation of a concept *C*, the *IX* current to all neurons in Ψ(*C*) is the same. Competition in the feature layer occurs because of the recurrent excitatory and inhibitory currents.

Therefore, after the presentation of a new concept *C*, once the first set of input spikes arrive at time *t*_1_, before any neuron spikes, all neurons in set Ψ(*C*) receive the same current. As a result, some time after *t*_1_, all neurons in Ψ(*C*) will fire at the same time - let us call the time when this happens *t*_*F*_. After *t*_*F*_, recurrent currents are active, and competition comes to play in the feature layer. After this, if no neurons fire till *t*_2_ (i.e. the time of the next input spike) then recurrent currents are not enough for sustained activity in the system. Therefore, objectives *O*1 and *O*2 will not be satisfied, and the system will not be operable. Else, the neurons that fire first after *t*_*F*_ would dominate the competition in the feature layer - as these neurons will reinforce their neighbors and inhibit non-neighbors - and influence which attractor is active at the end of the concept cycle. Therefore we consider this case in the following discussion.

Since we know exactly which neurons fired at *t*_*F*_, we can predict which are the first neurons that will fire after this and when. This is the basis of the simplified model. We use this model to predict what happens between *t*_*F*_ and the time of the first spike after *t*_*F*_ or *t*_2_, whichever is earlier.

We take *t*_*F*_ as the *time of reference* and denote *t*′ as the time elapsed since *t*_*F*_ (i.e. *t*′ = *t* − *t*_*F*_). We first calculate *IX, RX* and *RN* currents at *t*_*F*_. The decay of input kernel functions 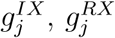 and 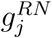 (equation 6) are modeled by exponential decay. Since neurons that receive *IN* currents do not participate in the activity during that concept cycle, they are not included in this discussion.

Following equation 6, the *IX* current at *t*_*F*_ depends on *t*_1_, the time at which the first input spike arrived:

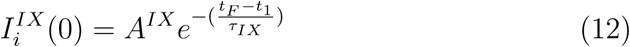

As can be seen in equations 6 recurrent currents are dependent on the neurons that are active in the excitatory receptive field of a neuron and its inhibitory receptive field respectively. So *RX* and *RN* currents at *t*_*F*_ are:

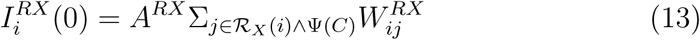

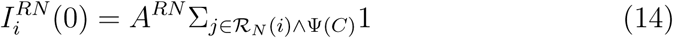

where ℛ_*X*_(*i*) is a set of all neurons in the excitatory receptive field of neuron *i*, and ℛ_*N*_ (*i*) is a set of all neurons outside its excitatory receptive field (and therefore all the neurons that it has inhibitory connections with). *C* is the current concept.

Before the first spike occurs after *t*_*F*_, the currents at time *t*′ are estimated to be

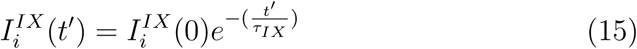

Similarly, for the recurrent currents,

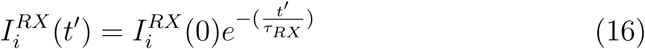

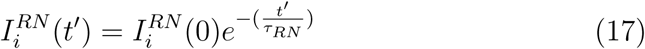

The currents calculated in the above equations drive the membrane voltage (equation 4). Also substituting these current equations in equation 4 and finding the integral, we can find a closed form equation for the membrane voltage as follows:

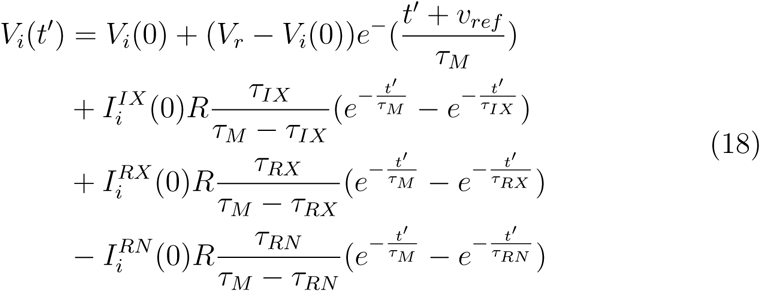

where *v*_*ref*_ is the refractory period of the system. The voltage and current equations above consist of the simplified model. This model can be used to analyze the system.

## 4. Objectives

To achieve the desired functionality, we identified two objectives that the system is required to fulfill:

(*O*1) Throughout the concept cycle, at least one neuron should be spiking. Since neurons do not spike in isolation due to the system’s architecture, some neurons should be spiking. Neurons should not all deactivate in the middle of a concept cycle.

(*O*2) For the initial context, the system should be able to identify categories, and create initial attractors corresponding to those.

In addition, there are two more objectives, which we hypothesized would improve the performance of the system.

For the next objective, we define a parameter, *Ψ* known as the *spread*. For each neuron *n* in an attractor, the spread of the neuron *Ψ*(*n*) is the number of neurons within the attractor outside its receptive field.

Within an attractor, the neuron (*n*_*max*_) with the maximum *Ψ*(*n*) is chosen, and so *Ψ*(*n*_*max*_) is known as the *spread of the attractor*. Among the initial attractors, the attractor with the largest *spread* is chosen to represent the *spread of the system* for that parameter combination.

When the attractors have a larger spread, we expect more interference between different attractors, and therefore, the system will perform less well. Further, when the attractors have a smaller spread, it will be easier to analyze the system.

(*O*3) The *spread* of the system should be less than *λ*.

Finally, if the new attractors are spatially separate from the initial attractors, there will not be interference between the initial and new attractors, and the performance would be better. Therefore, the fourth objective is:

(*O*4) Once initial attractors have been created and there is a context switch, the new attractors created should be spatially separate from the initial attractors. In other words, no part of the new attractor should fall within the receptive field of any neuron in the initial attractor.

For the rest of the paper, we will term the four objectives *O*1, *O*2, *O*3 and *O*4 respectively.

## 5. Results

### 5.1. Design Space Exploration

The RN currents were found to be very important for desired system dynamics. It was observed that when *τ*_*RN*_ was low, the objective *O*4 was not fulfilled and this hindered the system performance. On the other hand, when *τ*_*RN*_ was high, *O*4 was indeed fulfilled, and the performance was good. When *τ*_*RN*_ increases, there is an increase in the overall inhibitory current in the system. To maintain the system’s balance, it was noted that increasing *τ*_*RN*_ had to be accompanied by a corresponding increase in the excitatory current. There are two ways to increase the excitatory current contribution of a spike–increase the amplitude (*A*) or time constant (*τ*) of its corresponding kernel function. Increasing *τ*_*RX*_ or *A*^*RX*^ would lead to an increase in positive feedback in the system, and therefore lead to instability, and increasing *τ*_*IX*_ would not allow enough time for the attractors to learn - as the input comes into all neurons equally, competition in the network starts only after the input decays. Therefore, we chose to increase *A*^*IX*^ to complement the increase in *τ*_*RN*_.

The observation above leads to the following questions:

1. Why did the performance change with change in *τ*_*RN*_ ? What is the range of *τ*_*RN*_ for which *O*4 will be fulfilled?
2. How much excitatory current *A*^*IX*^ is required to balance the inhibitory current (*τ*_*RN*_)? What happens when either is too high relative to the other?

In order to answer these questions, we did a design space exploration of the system against objectives *O*1 − *O*4 to determine the parameter space of operation against these objectives. Further, the activity of the cerebral cortex is thought to depend on the precise relationship between synaptic excitation and inhibition [37]. Although this model is very simplified, it is based on principles that are commonly used to model the cortex. As a result, the experiments described in this section can qualitatively examine the ways in which this interaction takes place and provide directions for further research.

The system was tested on 600 concepts, from two contexts. The first 300 concepts belonged to the first context, and the next 300 concepts belonged to the second context. In these experiments the range of *A*^*IX*^ was from 15 nA to 110 nA with an interval of 5 nA, while the range of *τ*_*RN*_ was from 5 ms to 65 ms, with an interval of 5 ms. The system was tested at each combination of *A*^*IX*^ and *τ*_*RN*_ for Objectives *O*1 to *O*4.

#### 5.1.1. O1/O2 Experiments

##### Motivation

Both objectives *O*1 and *O*2 depend on the spiking activity of the neuron. By predicting whether or not there is some spike in the system, we can estimate whether *O*1 and *O*2 will be fulfilled. The experiment for doing so is described below.

##### Experiment

Initially weights are all small, so the neuron that receives most excitatory activity and least recurrent activity is constant for all parameter settings (and depends solely on the number of active neurons in a neuron’s excitatory receptive field). We chose that neuron (say 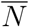) and checked if it was able to fire a spike after *t*_*F*_ but before *t*_2_. This was checked using both the simplified model and simulations.

A comparison of the predictions from the simplified model and simulations from the complete system were done to examine whether the neuron spiked - this is shown in Figure 5a. The results of *O*1 and *O*2 for all the values of *A*^*IX*^ and *τ*_*RN*_ are shown in figure 5b. Finally, one column in 5b is taken - where *τ*_*RN*_ = 15*ms* and the time taken for 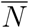 to spike after *t*_*F*_ was plotted for all values of *A*^*IX*^ and this is shown in 5c. In order to examine the increase in speed in the simplified model, we compared the time taken to perform the experiment in 5c by both the simulations and the simplified model. A comparison of the time taken to run the simulations from the start of the concept cycle till the firing of the first spike after *t*_*F*_ was observed and this was compared with the time taken to run the simplified model for the same.

**Figure 5:**
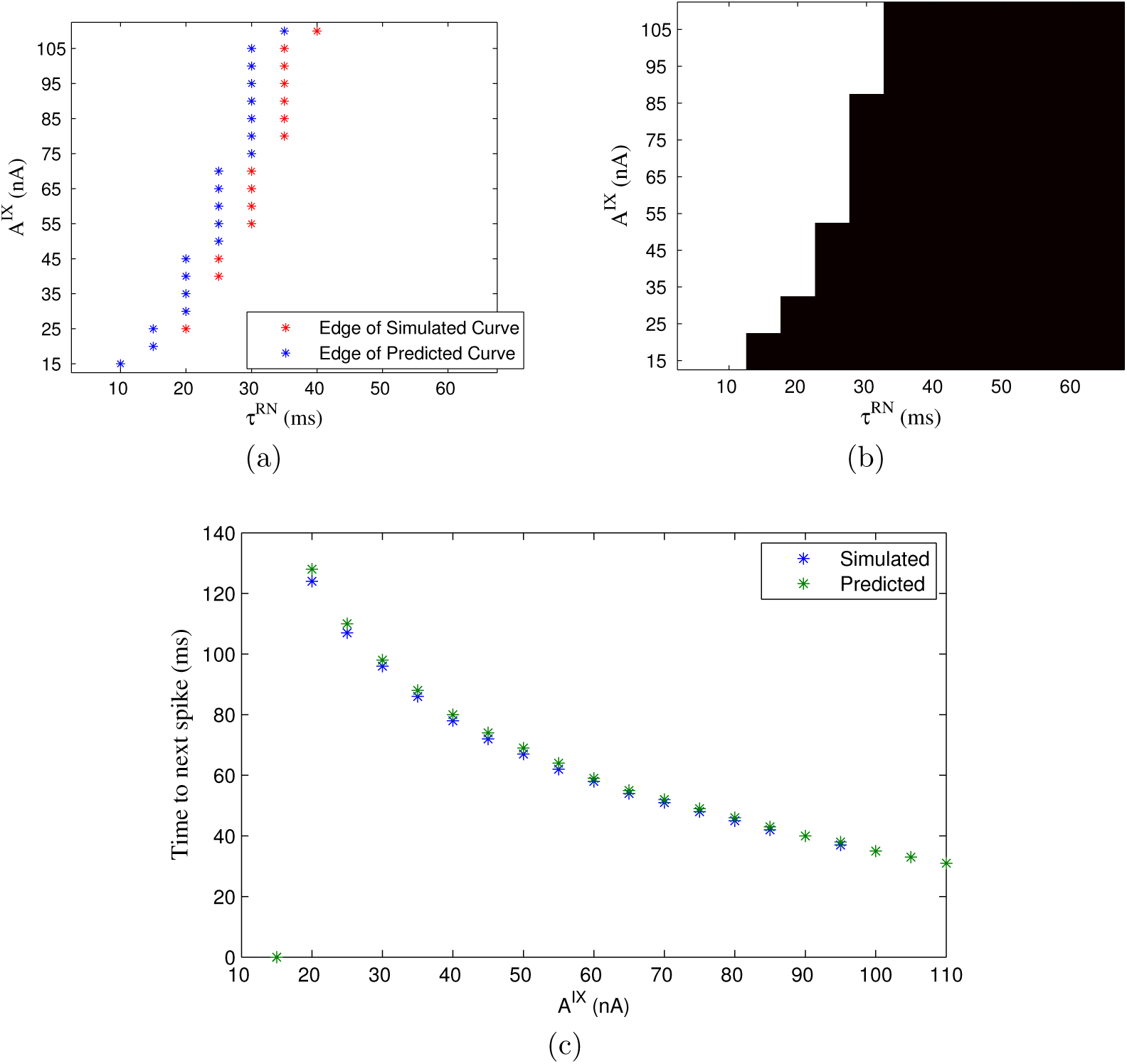
(a.) Predicted vs. simulation - will a neuron spike? Each set of symbols represents the boundary between a ‘Yes’ and a ‘No’ - the area to the left of the boundary is a ‘Yes’ and the area to the right is a ‘No’. See legend for details. (b.) Results of simulation for whether ‘O1’ and ‘O2’ are fulfilled. White color represents parameter values for which both objectives are fulfilled, and the black represents parameter values for which the objectives are not. (c.) Time of spike (how many ms after *t*_*F*_ does the first neuron spike?) when *τ*_*RN*_ = 15*ms* - Predictions vs. simulations. Time of 0 indicates no spike.

##### Results and Analysis

From Figure 5a it is seen that except for a few end points, the predictions match the simulations well. Averaging over all values of *A*^*IX*^ the simulations took 5.9425 seconds, while the simplified model took 0.0015 seconds showing that the simplified model made predictions about the actual model 3905 times faster on average for this case.

The above criterion is a necessary but not sufficient condition for both *O*1 and *O*2. Comparing figures 5a and 5b, it does give a good estimate of whether *O*1 and *O*2 conditions are met.

As seen from figure 5b, in the top left part of the graph, *O*1 and *O*2 conditions are met. The boundary follows a diagonal - indicating that for higher values of *τ*_*RN*_ higher values of *A*^*IX*^ are required to satisfy *O*1 and *O*2. The boundary indicates the maximum value that *τ*_*RN*_ can have for any value of *A*^*IX*^ so objectives *O*1 and *O*2 are fulfilled. As expected, when inhibition is too high relative to excitation, activity cannot be sustained in the network, and the required attractors cannot be created (see bottom right of figure 5b).

It is somewhat intuitive that with higher *A*^*IX*^, the neurons will get a higher voltage, and will spike, and lower *τ*_*RN*_, the neurons will not spike. Here we show how the simplified equation explains this. From equations 12 and 18 we can see how when *A*^*IX*^ is higher, any neuron is likelier to spike faster, as it gets more voltage.

For *τ*_*RN*_, we can rewrite equation 18 with just the *RN* component as follows:

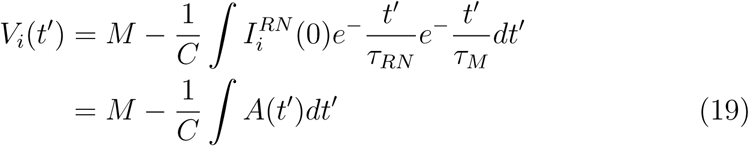

where all the other terms are referred to as *M* for simplicity. We can see that if −*A*(*t*′) is higher, then *V*_*i*_(*t*′) is higher and vice versa. Taking *log* of this term we get:

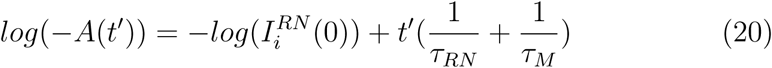

From equation 20 above, it is easy to see that the more *τ*_*RN*_, the lesser −*A*(*t*′), and the lesser *V*_*i*_(*t*′) will be (equation 19). Therefore an increase in *τ*_*RN*_ makes it harder for a neuron to spike.

#### 5.1.2. O3 Experiments

##### Motivation

As the objective *O*3 is dependent on the parameter *Ψ*, it is necessary to check this criterion for different values of *Ψ* and see how the results vary with the different values.

##### Experiment

For all values of *A*^*IX*^ and *τ*_*RN*_ and for different values of the parameter *Ψ*, the system was examined to see if objective *O*3 was satisfied. The simulation results are given in Figure 6.

**Figure 6:**
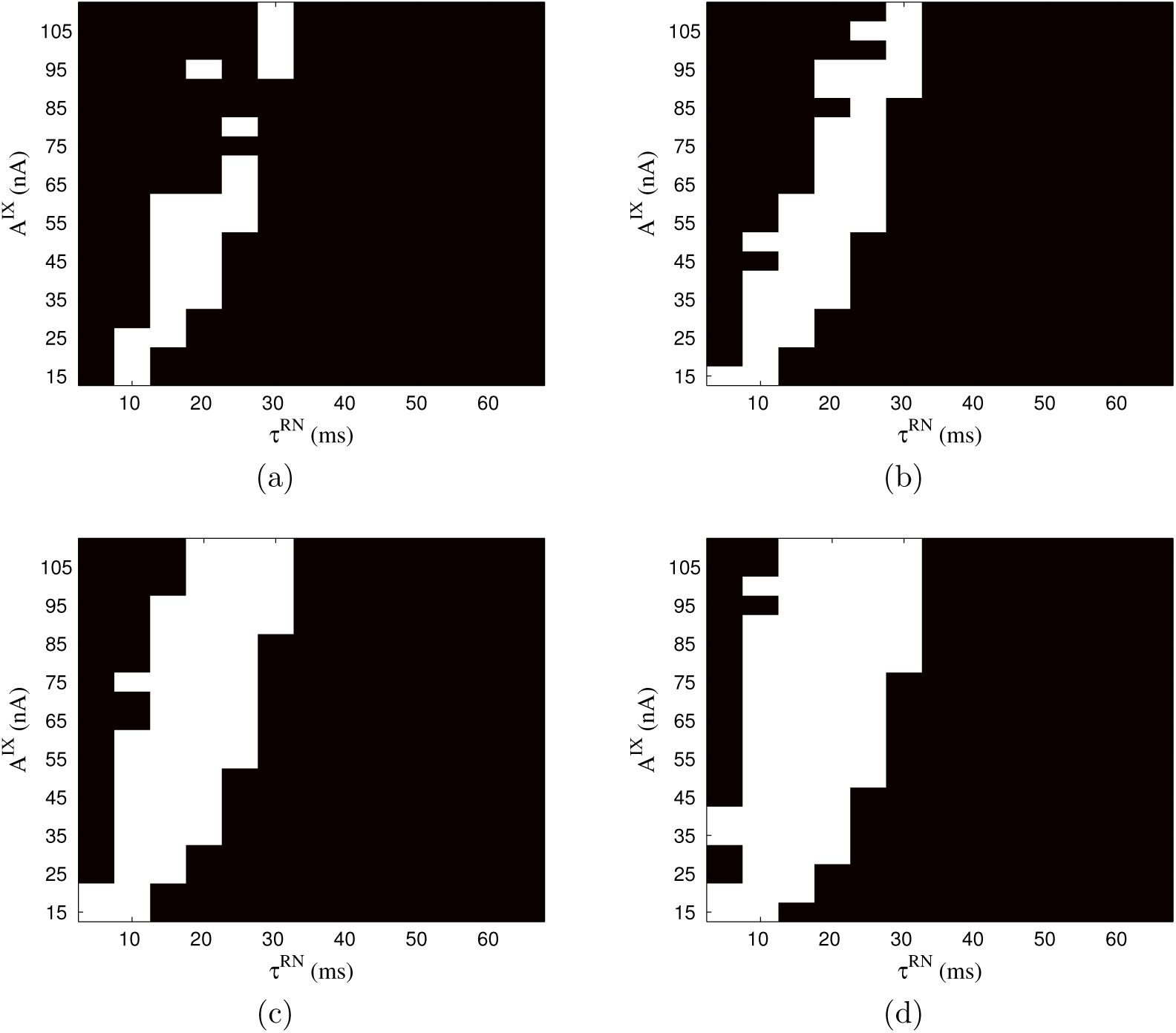
O3 Simulation results for *Ψ* = (a.) 12, (b.) 15, (c.) 20 and (d.) 25.

##### Results and Analysis

As can be seen, the values for which objective *O*3 is fulfilled follows a diagonal. On the right side of the diagonal, no attractor is formed, and on the left side of the diagonal the attractor is larger than the *Ψ* threshold. The diagonal gets larger as *Ψ* gets larger, with the units on the far left hand side alone being off, for the highest value of *Ψ*. This indicates that high values of *A*^*IX*^ coupled with low values of *τ*_*RN*_ result in larger attractors - the larger *A*^*IX*^ is compared to *τ*_*RN*_, the larger the spread of the attractors. The reason for this is explained as follows.

We analyzed the results with the simplified model. It was seen that neurons close to the centre of the attractor always got enough positive current (both from the input and recurrent neurons) for relatively low as well as high values of *A*^*IX*^. On the other hand, neurons further away got less positive current when *A*^*IX*^ was relatively low, and were not able to fire before inhibitory current from the already firing neurons took over, and prevented firing, resulting in smaller spread. However, when *A*^*IX*^ was high, neurons further away from the centre of the attractor were able to fire before inhibitory current was high enough to prevent them from doing so. Once they started firing, they had enough positive recurrent current from their excited neighbours, so they continued to be a part of the attractor. This is explained in figure 7. This is why higher *A*^*IX*^ without a corresponding increase in *τ*_*RN*_ results in larger attractors.

**Figure 7:**
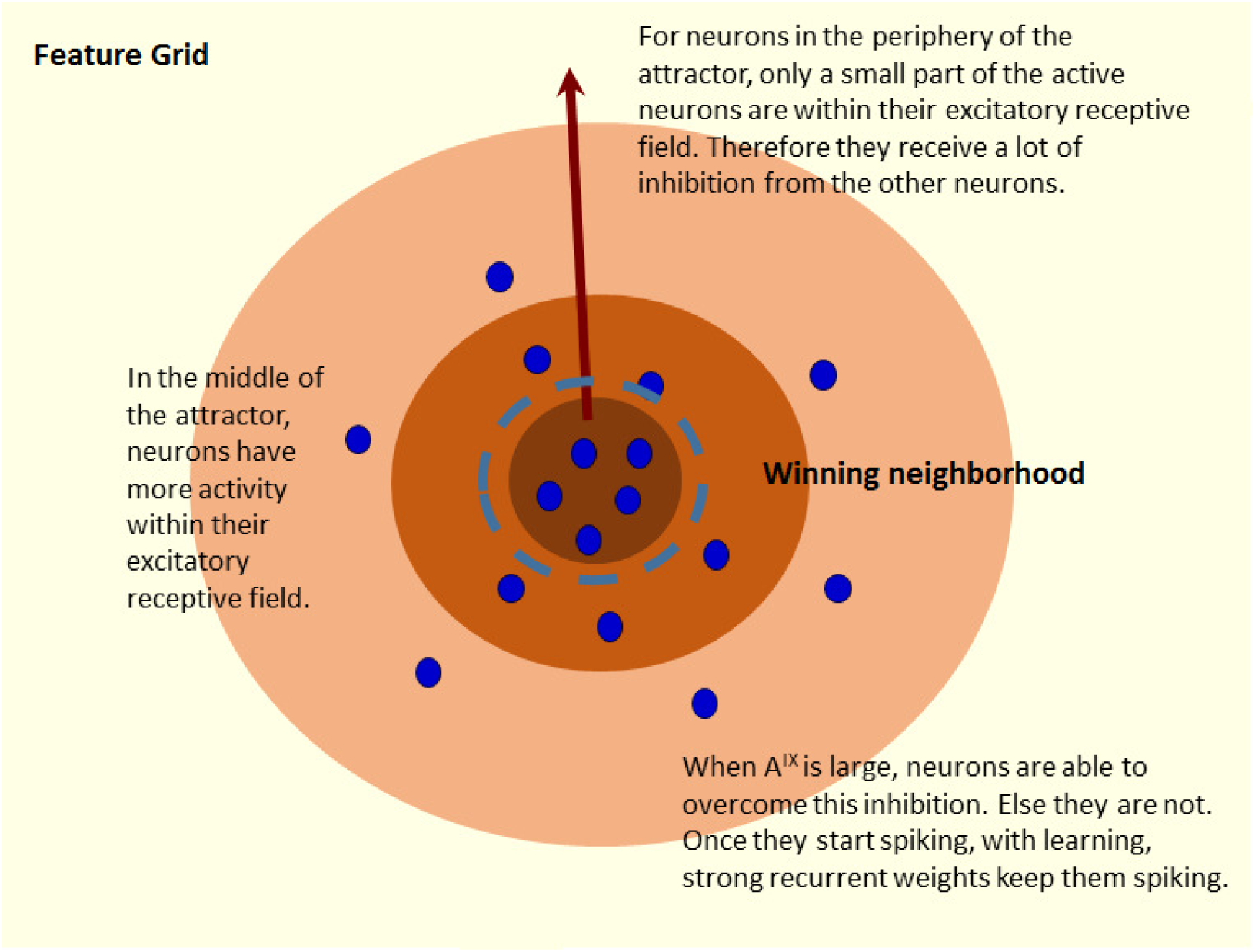
Explanation for *DSEO*3 results.

#### 5.1.3. O4 Experiments

##### Motivation

We now come back to the observation that was raised in the beginning of this section (section 5.1) - that when *τ*_*RN*_ was low (= 5 ms), *O*4 was not fulfilled, and when *τ*_*RN*_ was high (= 30 ms), *O*4 was fulfilled. We first use the simplified model to understand why this occurs, and analyze the parameter range in which *O*4 is likely to occur. At the end of this section, we compare it with the simulation results for *O*4.

We observe that under some conditions, after a context switch, when new concepts belonging to a new category are presented, parts of a previously learned attractor (belonging to a previous category) gets activated. This biases the competition towards the area around the initial attractor. As concepts keep getting presented, feature units that are part of the new category become more active and strongly connected, but these are close to the previous attractor due to the initial biasing of features of the old attractor (which naturally activate their own neighbors). Therefore, a new attractor is created overlapping or close to a previous attractor.

The obvious question is - under what conditions, or parameter values does this occur? We revert to the simplified model to answer this question. After a context switch, when a concept *containing features belonging to a previous category* is presented, either some previous attractor neurons are activated or a completely new attractor is formed. This depends on which neuron spiked first after *t*_*F*_ - as this is the step that strongly biases competition towards the winning attractor. We analyze which neuron is likely to spike first in different parameter settings. In this comparison spike rates do not matter, since *one* spike of all neurons at time *t*_*F*_ determines which neuron is going to spike next, which in turn determines which attractor will win.

We consider the magnitude and not signage of *RN* currents for ease of analysis. Since all neurons get the same *IX* current, the competition between two neurons is directed by the amount of *RX* and *RN* currents that the neurons receive. *RX* currents depend on the number of active neurons in a neuron’s receptive field as well as the excitatory weights between the neurons. Since neurons in the initial attractor are learned, their weights are relatively high compared to the weights of neurons which are not part of any attractor yet. So their *RX* currents are higher even if they have less neurons active in their receptive field. Any two neurons receive the same *RN* current from all active neurons in the system except for neurons in each other’s receptive field (each neuron will not get any inhibitory current from neurons in its own receptive field). The neuron with more units in its receptive field will receive less *RN* current than its competitor, since less neurons are providing inhibitory current. Therefore, the neuron with the *least RN* currents is the neuron with the maximum number of neighbours in its receptive field. Using the analysis above, we design the experiment as follows.

##### Experiment

After presenting the 100 initial concepts, there was a context switch. A concept was presented such that it shared some (but not all) few features that were a part of the initial category. For both high and low values of *τ*_*RN*_ (we chose *τ*_*RN*_ = 5 ms for *low* and *τ*_*RN*_ = 30 for *high*) we chose two neurons each for comparison, *N*1, an active neuron that is a part of the initial attractor and the current concept, and *N*2, the neuron with the maximum active neurons in its receptive field. Neuron *N*1 is expected to have higher *RX* current as it is unlikely for *N*2 to be a part of any attractor. Neuron *N*1 will have higher *RN* current than *N*2 as well (see how *N*2 is chosen). If *I*_*N*1_ is greater than *I*_*N*2_, *N*1 fires first, else *N*2 fires first. We take the simplified model, and use the equations to understand the conditions under which *N*2 fires first.

##### Results and Analysis

For neuron *N*2 to fire first, membrane potential *V*_*N*2_ should reach *V*_*th*_ before *V*_*N*1_. So using equation 4 and subtracting *V*_*N*2_ from *V*_*N*1_ we get:

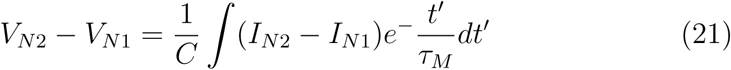

It is obvious from the equation that the more positive *I*_*N*2_ − *I*_*N*1_ is, the more positive the integral, and the LHS in equation 21 will be. Further, the difference *I*_*N*2_ − *I*_*N*1_ can be expressed in terms of the component currents, *IX, IN, RX* and *RN*. However, since *IX* and *IN* are the same for both the neurons they cancel out, so only the *RX* and *RN* components are left. *I*_*N*2_ − *I*_*N*1_ is expressed therefore as:

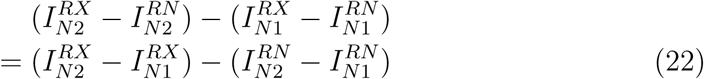

Now since we picked *N* 1 with the highest *RX* currents, 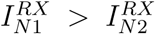, and since we picked *N* 2 with the least *RN* currents, 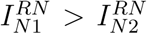. So both components in the final equation in 22 are negative. Rearranging this, and simplifying the equations based on 16 and 17, equation 22 will be more positive if:

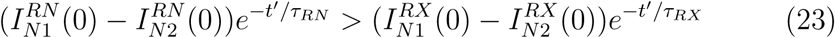

We denote difference between the currents with a subscript *diff* i.e. 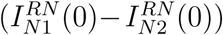 is expressed as 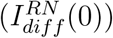 and similarly for *RX* currents. We take *log* of the equations and we get:

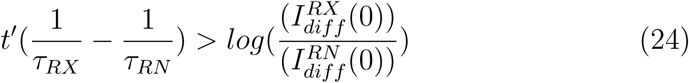

As is obvious from the equation above, the higher *τ*_*RN*_, the more likely the equation is going to be satisfied. Also, while we cannot guarantee that if *I*_*N*2_ *> I*_*N*1_ after some specific time, the neuron will spike first, since it involves the integral, we can say that the more positive the difference between *I*_*N*1_ and *I*_*N*2_, the more likely that *N* 2 spikes first. From the equation 24 therefore can state two things: (1.) For higher values of *τ*_*RN*_, it is likely that *O*4 is fulfilled. We can expect 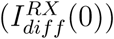 to be higher than 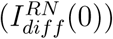 as the former is the difference between the weights of a learned and unlearned attractor respectively, while the latter is just due to the number of neighbors that are active in the excitatory receptive field of each neuron (and therefore due to chance). However, with a large enough *τ*_*RN*_ this difference can be overcome. (2.) The larger *A*^*IX*^ is, the larger 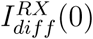 is, (see equation 16) so we need higher values of *τ*_*RN*_ for equation 24 to be satisfied. We see the results of *O*4 in figure 9, and we see that both statements made above are indeed satisfied.

From this analysis a natural prediction arises - for relatively high values of *τ*_*RN*_ the neuron with the maximum number of neighbors in the receptive field will be the first to activate after *t*_*F*_ and the time to activation of neurons will follow the number of active neighbors in their receptive field. Figure 8c shows that this prediction is indeed satisfied - the rank of the time to spike of neurons follows the number of neurons in their excitatory receptive field.

**Figure 8:**
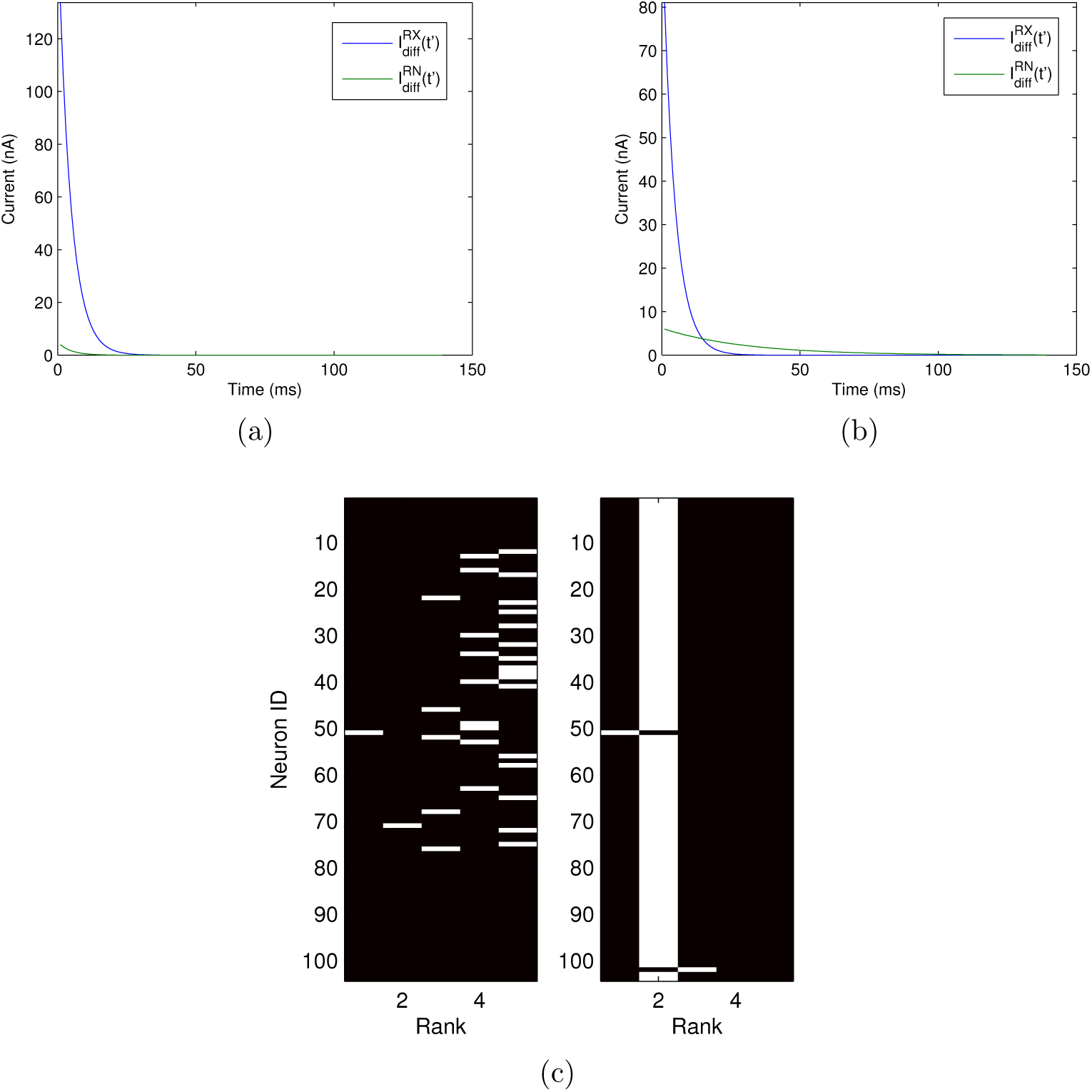
Above: Difference in currents as shown in 24 respectively for *τ*_*RN*_ = 5 ms (left) and *τ*_*RN*_ = 30 ms (right). As can be seen, when *τ*_*RN*_ = 30 ms although the difference in *RX* currents starts high, the difference between *RN* currents eventually exceeds it. This is predicted in the explanation following equation 24. Below: Neurons with the maximum neighbors were predicted to spike first after *t*_*F*_. Their predicted rank and actual rank (ranked according to time of spike) are compared. For ease of visualization, only those neurons that fall into the first 5 ranks are shown in the figure.

**Figure 9:**
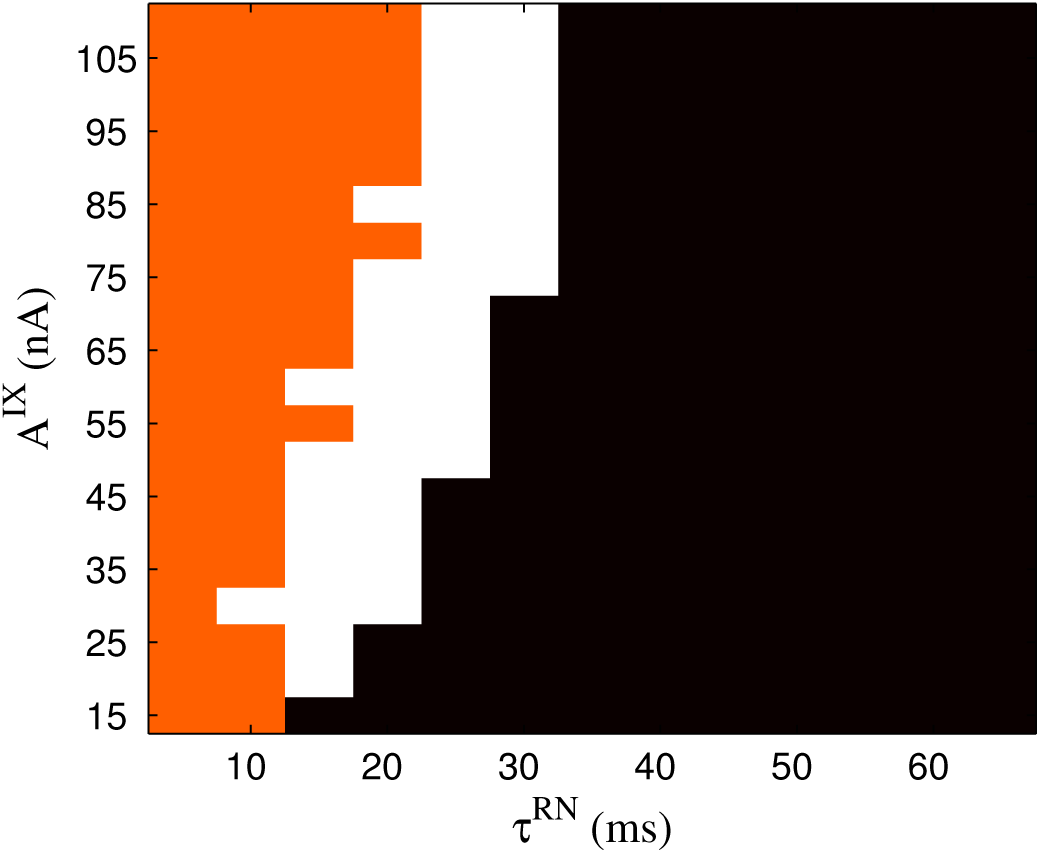
O4 simulation results. Orange indicates that O4 is not fulfilled, white indicates that O4 is fulfilled, and black indicates that the attractors are not created at all.

#### 5.1.4. DSE Simulation Results

Finally we performed design space exploration for the system, and checked the performance for different values of the parameters - the results are given in 10. The system was trained for each combination of *A*^*IX*^ and *τ*_*RN*_ values described above. During testing, the attractors are already learned and the objective is activate learned attractors, and not other neurons. Hence, for all resting, *A*^*IX*^ was set low, to 15 nA, and *τ*_*RN*_ was set as high as possible so as to not prevent activity in the system - we chose 10 nA. During testing, the units that have greater than a spike rate of *θ* for the final *k*% of the training time are considered the output of the system. *Performance error* is calculated as the sum of # of false positives + # of false negatives. As the total number of concepts × total number of possible features in a concept is the same, the performance error in different cases can be compared without normalizing. If performance error is less than *e* (= 15 in our simulations), the performance is defined to be acceptable, else it is not. The simulation results are shown in figure 10.

**Figure 10:**
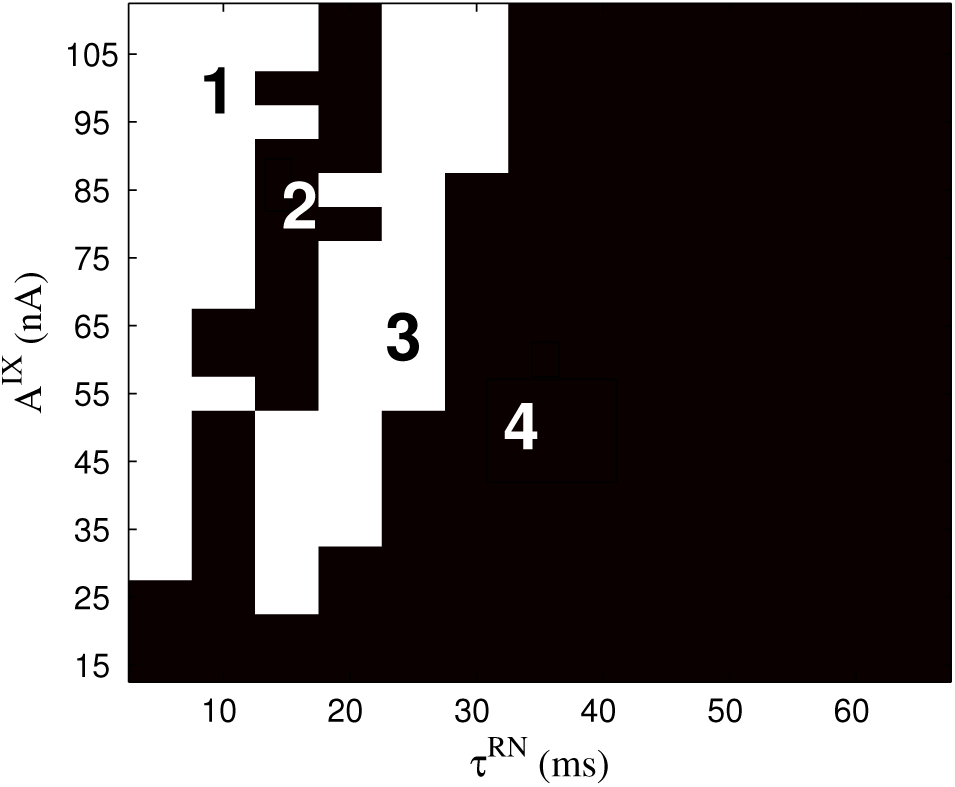
Final results over the parameter space for *A*^*IX*^ and *τ*_*RN*_.

We examined the reason for the performance given in the figure. We confirmed that *O*1 and *O*2 were necessary for the functionality, and examined whether *O*3 and *O*4 were necessary, and the role they played in the functionality. As can be seen in the figure the range of parameters has been divided into four based on performance - these are labelled part 1, 2, 3 and 4 in the figure - these have been examined separately. In part 1 and 3, the system does exhibit desired functionality, while in part 2 and 4, it does not.

Part 4 has bad performance because *O*1 and *O*2 are not satisfied (compare with figure 5b). Part 3 has good performance as *O*1, *O*2 and *O*4 are satisfied (compare with figures 5b and 9). Separate attractors corresponding to different categories are created and they do not interfere with one another. In Parts 1 and 2, *O*1 and *O*2 are satisfied but *O*4 is not (compare with figure 5b and 9). Then how does part 1 give good performance but part 2 does not? In Part 1, the spread of the attractors is relatively high (check figure 6c). This is considerably larger than a neuron’s receptive field. So neurons with excitatory connections and inhibitory connections will be a part of both the attractors. In this case, although two attractors exist in the same spatial area, when the input activates the more dominant attractor, there is inhibitory current preventing the other attractor from getting activated. As a result, the performance is good in part 1. In part 2, the spread of the attractors is small, and there is no inhibition between the two attractors. So when features of the dominant attractor, and some features of the second attractor are active both the dominant attractor and parts of the second attractor will become active. As a result, the performance will be compromised.

This leads to a surprising, but interesting result - when *O*3 and *O*4 are both not fulfilled the system’s performance is good. However, if *O*4 alone is not fulfilled, the system’s performance is not good.

### 5.2. Comparison with biphasic STDP

As mentioned earlier, we chose to train the system with triphasic STDP learning rule because it makes intuitive sense to have the neurons connected to one another through symmetric weights - the weights between two neurons does not imply causality (i.e. B spikes as a result of a spike in A) but correlation (feature B and A are highly correlated with one another). If two features A and B are highly correlated to one another, the connections between A and B, as well as B and A should be high, and vice versa. Therefore, it was not reasonable to use a learning rule that increased weights in one direction, but decreased them in the other direction.

The system with the learning rule described above was run over the entire parameter space and the results are shown in figure 11.

**Figure 11:**
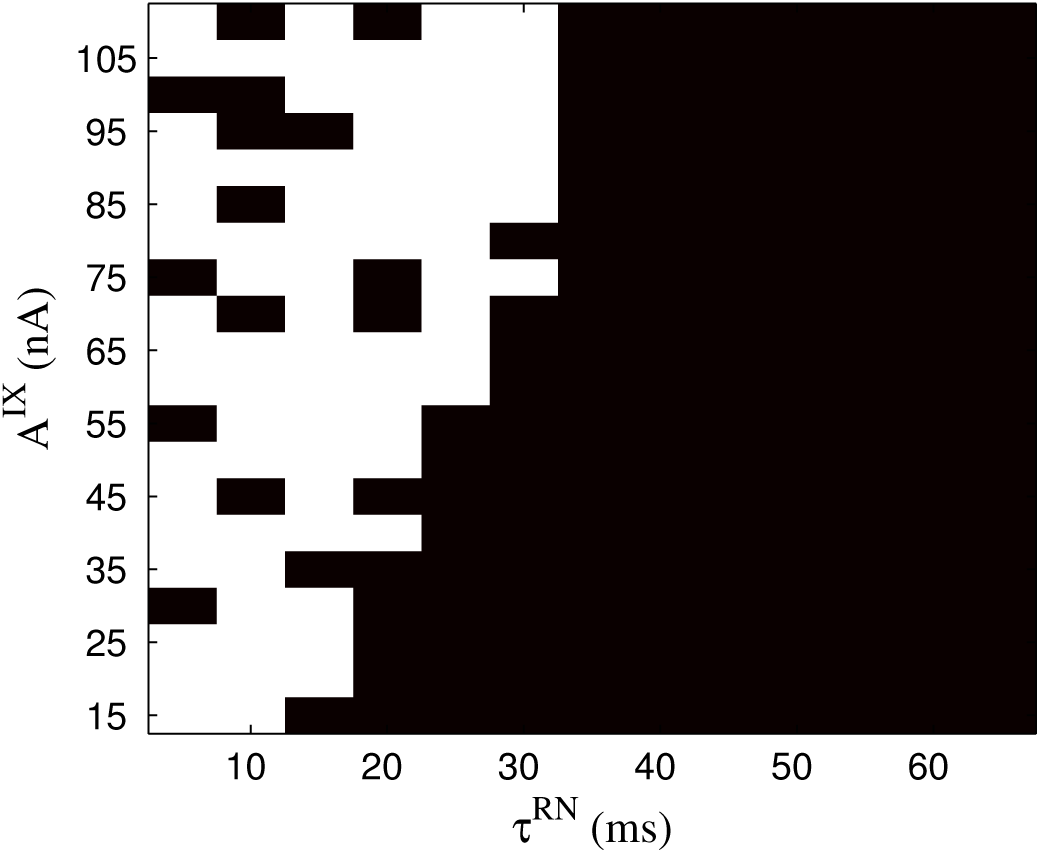
Final Results with biphasic STDP over the parameter space for *A*^*IX*^ and *τ*_*RN*_

#### Motivation

We also compared the time it took to train the system with both learning rules. The protocol for the experiment is given below.

#### Experiment

All weights in the system are subject to decay until they are above the *stabilization threshold*. Therefore an attractor is considered *stable* if all the weights between neurons in the attractor have crossed the *stabilization threshold*. In this experiment we calculate the number of concepts required to create a stable attractor. We use the term Γ_*c*_ to denote the fraction of the entire attractor that is created at the *c*th concept, and is calculated as:

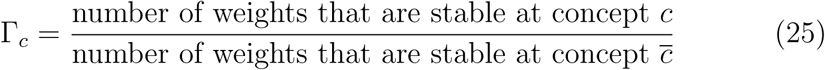

where 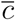 is an extremely large number. The idea is that when the system is trained for an indefinitely long time, the attractor is stable, and there will be no further changes. This is the *ideal* attractor, and Γ_*c*_ measures the deviation from the ideal attractor. Here 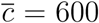 as all attractors are stable much faster than that.

The number of concepts the system needed to be trained on before Γ_*c*_ fell below *γ*(= 0.05) was calculated for each combination of *A*^*IX*^ and *τ*_*RN*_. This was repeated four times each using four different sets of concepts generated in a similar manner, and was done for both triphasic and biphasic STDP learning rules. Average values were taken for each parameter combination. The results are shown in figure 12.

**Figure 12:**
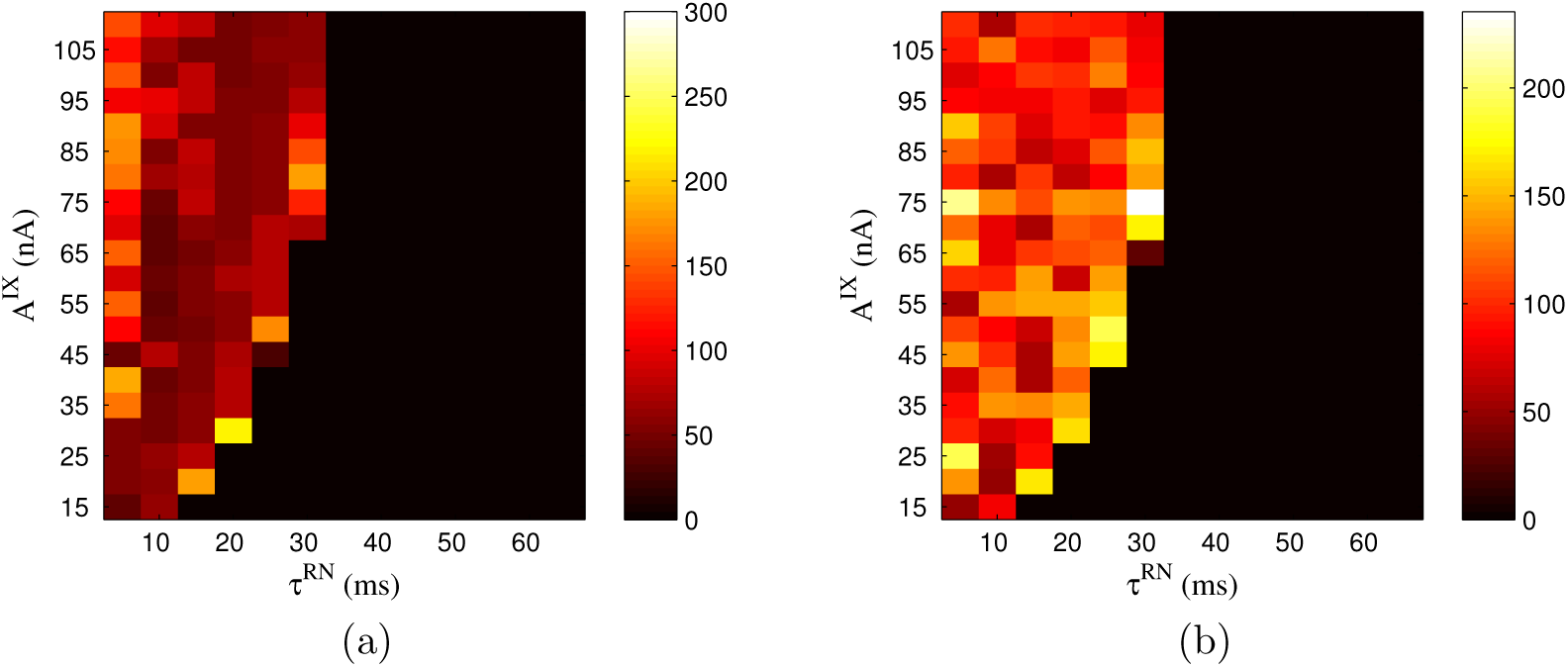
Number of concepts that the system had to be trained on before Γ_*c*_ fell below *γ*(= 0.05) for (a.) triphasic STDP and (b.) biphasic STDP, averaged over 4 runs.

#### Results and Analysis

In these results, we notice that although there is not a considerable degradation in the performance of the system, there is a large difference in the training time for these two cases. This is due to the nature of the learning rule. Using biphasic STDP learning rule, when two neurons (say *A* and *B*) spike temporally chose to one another (say, *B* spikes right after *A*), the weights between *A* and *B* increase but weights between *B* and *A* decrease. In a sequence of spikes (such as *ABABAB*) there is a net overall increase in the weights - the increase in the weights between the neurons overcomes the decrease in the weights - this is how attractors are created. However, with the triphasic STDP learning rule, whenever *B* spikes after *A* or vice versa, there is a symmetrical increase in weights in both directions. This is shown in figure 13. Triphasic STDP learning rule is resulting in exactly the characteristic that we need: a symmetrical increase in weights every time the spikes occur in close temporal succession. As a result, the weight increase occurs quickly, and the training time is less, given other parameters are comparable.

**Figure 13:**
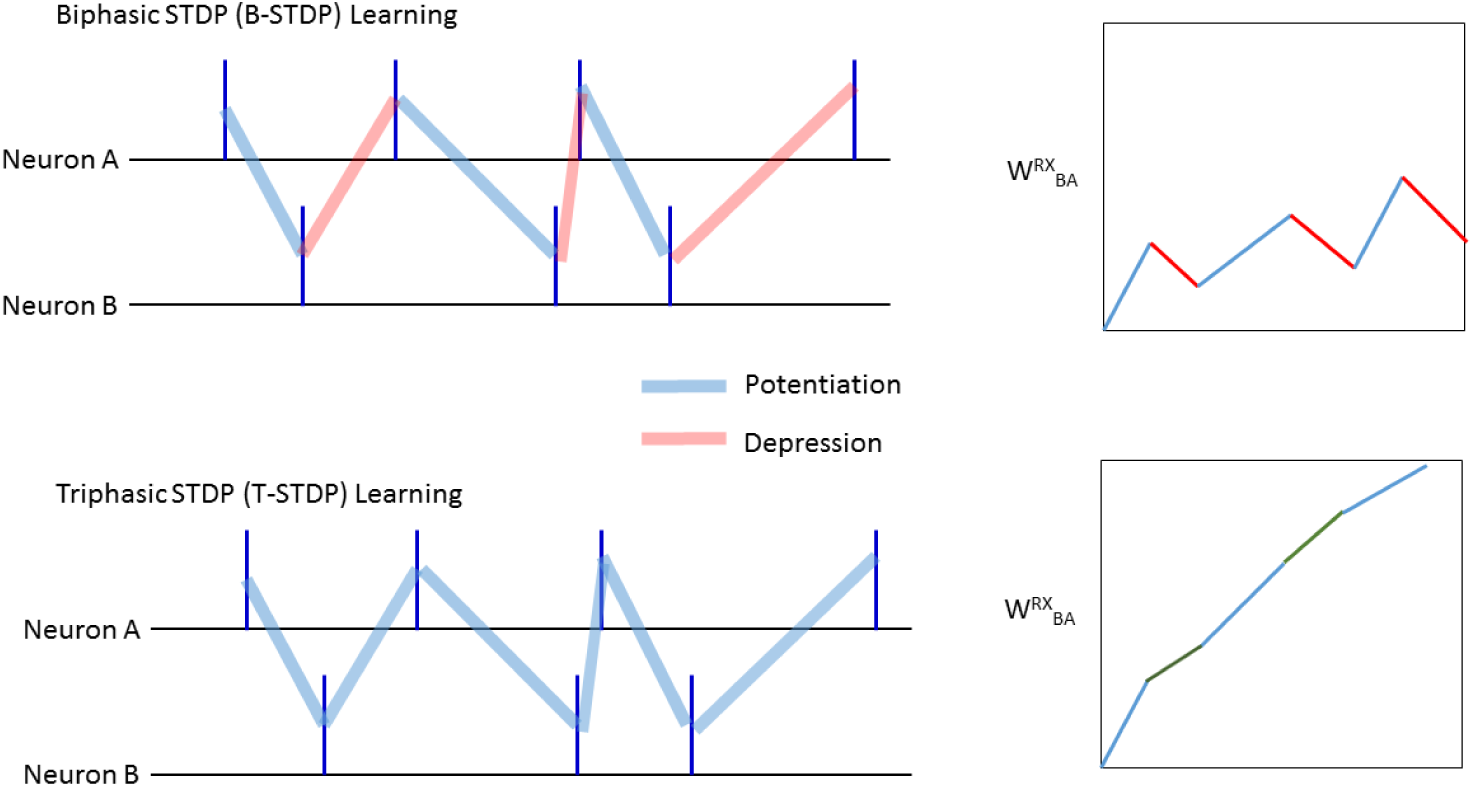
Top left: Spike raster for neurons *A* and *B* and learning in *B* − *STDP*. Top right: The resultant weight change. Bottom left: Spike raster for neurons *A* and *B* and learning in T-STDP. Bottom right: The resultant weight change. When two neurons (A and B) spike temporally close to one another, in B-STDP, there is potentiation and depression in the weights. In T-STDP there is only potentiation (depression occurs when neurons spike happen further apart - see 3).

Another thing we note in the results is that although the number of cases (i.e. *A*^*IX*^/*τ*_*RN*_ combinations) in which the system performs well is comparable in biphasic and triphasic STDP learning rules, the patterns of good performance are very different (see figures 10 and 11). In triphasic STDP, we clearly identified four parts in the parameter space, and analysed their performance. In biphasic STDP, the results are discontinuous. The explanation for this is given below.

In earlier results with the triphasic STDP learning rule, we observed that when *A*^*IX*^ was high compared to *τ*_*RN*_, large attractors were created (see figure 6). With biphasic STDP learning rule, however, in this case, we note something different - an attractor consists of several small disjoint parts, which are spatially separated from one another (i.e. no neuron in one disjoint part is excitatorily connected to any neuron in the other attractor). It is well established that symmetric learning rules in recurrent neural networks result in single attractors [38, 39, 40], while such guarantee cannot be made with asymmetric networks.

Additionally when *τ*_*RN*_ is higher, the attractors have a very small spread, and may not cover all salient features, resulting in incomplete attractors. Disjoint attractors and incomplete attractors give rise to more complex phenomena that are out of the scope of this paper. Discontinuities occur due to these phenomena.

## 6. Choice of Parameters

All of the parameters are based on biologically plausible ranges [41]. Most of the standard parameter values were chosen from a biologically plausible range of values, without other considerations, and these include *v*_*ref*_, *τ*_*M*_ and *τ*_*RX*_. *τ*_*RN*_ was initially set to be equal to *τ*_*RX*_ but different values of the parameter were explored in the DSE experiments.

In the system, *RN* currents ensure that any neuron that is not getting direct input from its input feature, but getting recurrent currents alone is not allowed to be active. Hence, *A*^*IN*^ and *τ*_*IN*_ were chosen so that the negative currents would override any positive current that was input into the neuron due to recurrent activity alone.

The range of EPSC amplitudes required, given the other LIF parameters, to ensure good spiking activity (activity was not so low that no neuron spiked, or so high that the neurons were spiking at the maximum spike rate, given the weights) was considered, and *A*^*RX*^ was chosen based on this consideration. *A*^*IX*^ was initially set to be equal to *A*^*RX*^ but different values of the parameter was explored in the DSE experiments.

*τ*_*IX*_ was set such that the *IX* currents will serve as a ‘booster’ current to allow activity in the system before recurrent currents take over, yet, allow for competition within the concept cycle. If *τ*_*IX*_ is too low, given the current parameters, there may not be enough activity in the system to sustain recurrent activity during the concept cycle. If *τ*_*IX*_ is too high, the input currents will act for too long, thus not allowing enough time for competition between the regions, to ensure that one attractor emerges during the concept cycle. *τ*_*IX*_ value was chosen from ballpark ranges that would satisfy these conditions and the parameter was not tuned.

*V*_*th*_ and *A*^*RN*^ were set such that activity is limited to one region, and not all units get inactive, given the other parameters. If *V*_*th*_ or *A*^*RN*^ was too high, not enough neurons would be active, given the other parameter values, and if *V*_*th*_ or *A*^*RN*^ was too low, too many neurons will be active in more than one region. Parameter values were chosen from ballpark ranges that would satisfy these conditions, and the parameters were not tuned.

As can be seen, a balance between excitatory current and inhibitory current is necessary for the proper functioning of such a system. This is one of the main conclusions of the DSE experiments as well. Hence after keeping some parameters stable, other parameters were chosen such that this balance would be maintained.

## 7. Discussion and conclusion

A summary of the system features and main results are as follows:

- An SNN model of category learning was created where categories are a combination of features.
- The system architecture is a hybrid between recurrent neural networks and self organized feature maps, and has been used commonly in other models. However, we use the architecture differently to model higher level category learning, and similar features are not necessarily topologically close. This is to facilitate the creation of diverse attractors.
- The system exhibits desired functionality for a range of parameters. A design space exploration of the system was conducted to examine this range.
- Four objectives were identified, and we examined whether the system needed to fulfill them to exhibit the desired functionality, and the range in which the functionality was exhibited. The analysis done for the system is an analysis for local excitatory-inhibitory networks [42] in general.
- We noted that for different ranges of parameters the system exhibited different behaviors.
- A comparison of T-STDP and B-STDP showed that T-STDP was in general better suited for this system. Although the performance was comparable, the training time was higher for B-STDP.

A very interesting result from the system is that the system exhibits good performance when the attractors are small, and are not overlapping (as expected), but when the attractors are larger and have inhibitory currents against one another, they can be overlapping and yet exhibit good performance. This is due to inhibitory currents inhibiting the less dominant attractor from getting activated. Thus we see that in one system with the same architecture, with the change of one parameter (i.e. the spread of the attractor) the criteria for good performance become completely different (i.e. overlapping attractors cause bad performance in one case and not the other). Evidently the brain uses such mechanisms - as it has similar operating principles and the same architecture throughout, but with some change in parameters, very different cognitive states are achieved. In particular, the behavior shown in this system may provide light to further research, to examine whether smaller attractors that have only positive connections can co-exist, while larger attractors compete and impede one another. In general, uncovering such mechanisms would provide fodder for further research into the brain’s operative mechanisms. This is an important step in banishing the homonculus.

One of the benefits of lateral excitatory and inhibitory connectivity cited by Mikkulainen is that excitatory and inhibitory connections play different roles in the cortex 1. As we saw in section 5.1.3, when *τ*_*RN*_ is low, the neuron with the largest *RX* current wins. However, if *τ*_*RN*_ is high, the neuron with the smallest *RN* currents win. This is because excitatory weights are potentiated during learning while inhibitory weights are left unchanged. As a result of this different learning, excitatory currents enable the most well learnt neurons to win, while inhibitory currents enable the neuron that has the most number of active neighbors to win. Again, as excitatory and inhibitory currents play different roles, there are two competing mechanisms, and this disallows the dominance of any one attractor. Further, once again, with a change of one parameter, *τ*_*RN*_, the system exhibits very different characteristics and performance. From the engineering viewpoint, when there are two separate mechanisms, i.e. excitatory currents and inhibitory currents, the complexity is being increased, therefore it is best that they play different roles.

Finally, we did a comparison with B-STDP and T-STDP, and showed that T-STDP is faster. Standard B-STDP implies causality, and certainly, in many of the brain’s mechanisms, causality is required. However, the detection of feature correlations requires the detection of coactivity rather than causality, and this is why T-STDP, a symmetric learning rule performs better than standard STDP. T-STDP is one such rule, but there may be other mechanisms. This needs to be explored to understand how the brain deals with coactivity in an optimal manner.

We chose recurrent connectivity as opposed to feedforward connections as we felt that recurrence captures coactivation rather than causality. In future work, a more rigorous comparison of feedforward and recurrent connectivities will be done.

## 8. Acknowledgement

This work was supported by MOE through grant MOE 2013-T2-2-017.

